# Decreased spinal inhibition leads to undiversified locomotor patterns

**DOI:** 10.1101/2022.04.21.489087

**Authors:** Myriam Lauren de Graaf, Heiko Wagner, Luis Mochizuki, Charlotte Le Mouel

## Abstract

During walking and running, animals display rich and coordinated motor patterns that are generated and controlled within the central nervous system. Previous computational and experimental results suggest that the balance between excitation and inhibition in neural circuits may be critical for generating such structured motor patterns. In this paper, we explore the influence of this balance on the ability of a reservoir computing artificial neural network to learn human locomotor patterns, using mean-field theory and simulations. We created networks with varying neuron numbers, connection percentages and connection strengths for the excitatory and inhibitory neuron populations, and introduced the anatomical *imbalance* that quantifies the overall effect of the two populations. We trained the networks to reproduce muscle activation patterns derived from human recordings and evaluated their performance. Our results indicate that network dynamics and performance depend critically on the anatomical imbalance in the network. Excitation-dominated networks lead to saturated firing rates, thereby reducing the firing rate heterogeneity and leading to muscle coactivation and inflexible motor patterns. Inhibition-dominated networks, on the other hand, perform well, displaying balanced input to the neurons and sufficient heterogeneity in the neuron firing rate patterns. This suggests that motor pattern generation may be robust to increased inhibition but not increased excitation in neural networks.

## 1 Introduction

Humans and other animals create rich and co-ordinated motor patterns during walking and running. With increasing locomotor speed, the stepping frequency increases and the pattern of muscle contraction changes, both in terms of the amplitude of the muscle contractions as well as their timing within the step cycle (Ivanenko et al., 2004, 2006; Lacquaniti et al., 2012). The muscle activation patterns are generated by locomotor central pattern generators (CPGs) within the spinal cord (Delcomyn, 1980; Ijspeert, 2008; Grillner and Wallén, 1985; Grillner et al., 1998). The spinal cord is not only capable of generating the rhythm of locomotion, but also of transforming this simple rhythm into the complex muscle contraction patterns observed during locomotion. In this study, we investigate how this pattern formation is influenced by spinal anatomical characteristics, specifically the excitation-to-inhibition-ratio.

The role of inhibition in CPGs has already been studied experimentally by blocking inhibition in animal models. When inhibition is blocked, the generated rhythm is undisturbed, but the flexors and extensors are active simultaneously and vary in phase (Cowley and Schmidt, 1995; Lanuza et al., 2004; Talpalar et al., 2013; Cohen and Harris-Warrick, 1984). This indicates that inhibition is essential for generating the alternating pattern between flexion and extension (Zhang et al., 2014; Britz et al., 2015). Similarly, inhibitory neurons have also been shown to play an important role in alternation between the left and right sides of the body (Lanuza et al., 2004; Talpalar et al., 2013; Kiehn, 2006; Jankowska, 2008; Quinlan and Kiehn, 2007). Inhibition thus plays a critical role in pattern formation of CPGs, but the mechanism through which it affects motor patterns is unknown. Therefore, we aim to study how the excitatory-to-inhibitory ratio affects locomotor pattern formation.

Separate spinal circuits have been identified that regulate rhythm generation on the one hand and pattern formation on the other. Rhythm-generating neurons are located medially and project locally, whereas pattern-forming neurons are located laterally and project to the lateral edge of the spinal cord, where the motoneuron pools are situated (Griener et al., 2013). Assuming separate rhythm generation and pattern formation circuits in modelling approaches can successfully replicate experimental phenomena such as non-resetting deletions (Lafreniere-Roula and McCrea, 2005), i.e. the finding that rhythm is maintained even when parts of the movement cycle are skipped (Rybak et al., 2006; McCrea and Rybak, 2007, 2008), as well as the effects of sensory stimulation on locomotor patterns (Rybak et al., 2006; McCrea and Rybak, 2007).

Rhythm generation by CPGs is most often conceptually modelled using the half-centre model (explained in Latash, 2012). This model assumes reciprocal inhibition between two neural pools (the half-centres), with one exciting extensor motoneurons and the other flexor motoneurons (Jankowska et al., 1967). When the extensor pool is active, it inhibits the flexor pool. After a while, the activity in the extensor pool decreases by some self-inhibitory process, such as fatigue (Brown, 1911; McCrea and Rybak, 2008; Latash, 2012). This releases the flexor pool from inhibition, allowing it to activate and concurrently inhibit the extensor pool until it, in turn, fatigues. The result is a rhythm of alternating activity between the flexor and extensor pools.

Pattern formation is necessary to extend the basic rhythm of alternating flexion and extension into the coordinated multi-joint muscle activation patterns that underlie locomotion. Muscle synergy analysis has identified a set of 4 to 5 invariant activation patterns – the synergies – that describe lower limb muscle activation during healthy locomotion (Dominici et al., 2011; Ivanenko et al., 2004; Chvatal and Ting, 2012; Clark et al., 2010; Gui and Zhang, 2016; Barroso et al., 2014). Such synergies can recreate locomotion in musculoskeletal models at a wide range of speeds (Neptune et al., 2009; McGowan et al., 2010; Aoi et al., 2019; Di Russo et al., 2023), but the mechanisms underlying their formation remain unclear. Conceptual half-centre models struggle to reproduce these diverse activation patterns. Some models propose coupled oscillators for each joint (unit burst generators, Grillner, 1981), or two half-centres acting at each joint (Li et al., 2017), which allows hip, knee and ankle extensors to vary slightly out of phase. However, these models still assume a strict alternation between flexors and extensors, thus failing to capture coactivation and the more complex activations of bi-articular muscles (Markin et al., 2012). Later models, therefore, extended the pattern formation networks to also account for bi-articular muscles (Shevtsova et al., 2016).

However, none of these conceptual models explain the concurrent inhibitory and excitatory neural activity observed in nature. According to the half-centre models, the input to a neuron in a given pool should be excitatory when that neuron’s pool is active, and inhibitory when the opposite pool is active. In contrast, intracellular recordings of spinal neurons show that motorneurons receive simultaneous excitatory and inhibitory inputs when their pool is active, rather than alternating excitation and inhibition (Berg et al., 2007; Petersen et al., 2014). A hallmark of this balance between excitatory and inhibitory inputs (EI balance) is a firing rate distribution across the neural population which is positively skewed (Petersen and Berg, 2016). These skewed firing rates cannot be recreated by reciprocal inhibition, as reciprocal inhibition leads to disinhibition of the active half-centre and thus to run-away activity (Berg et al., 2019). Recurrent inhibition must thus be present in spinal neural circuits. Since the half-centre models do not incorporate recurrent inhibition, they are unable to investigate its effects, thus indicating the need for an alternative model.

To further investigate the role of inhibition in pattern formation, we propose to use recurrent neural networks. Simulation studies have shown that such networks are capable of producing complex, high-dimensional signals, even in the absence of input or with a simple input signal (Funahashi and Nakamura, 1993). In reservoir computing (Verstraeten et al., 2007), a subtype of recurrent neural networks, linear readouts of the network’s activity are trained to reproduce a variety of target signals, while the internal connection weights remain static. Reservoir computing has been used to model how biological neural networks achieve a variety of tasks (Hinaut and Dominey, 2013; Bostrõm et al., 2013; Wyffels and Schrauwen, 2009). Moreover, it has proven highly effective in reproducing features of human locomotion such as the trajectories of markers placed on the human body (Sussillo and Abbott, 2009) and joint angles (Wyffels and Schrauwen, 2009; Hoellinger et al., 2013).

An additional advantage of using artificial neural network models is that their behaviour can be analyzed via mean-field theory (MFT), which describes the behaviour of large, complex systems in terms of their average effects rather than each of their individual components. MFT has been applied to investigate neural network dynamics, e.g., around the transition to chaos (Kadmon and Sompolinsky, 2015) or in networks with balanced excitation and inhibition (Harish and Hansel, 2015). Recurrent neural networks thus provide a versatile and insightful framework for studying neural pattern formation (Wyffels and Schrauwen, 2009; Hoellinger et al., 2013).

By modelling a network with recurrently connected neurons that mimic the excitatory and inhibitory interactions observed in spinal networks, we can investigate the importance of recurrent inhibition in the generation and modulation of motor patterns. While the functional EI balance in biological and artificial neural networks has been extensively studied (reviewed in, e.g., Hennequin et al., 2017; Sadeh and Clopath, 2021; Herstel and Wierenga, 2021; Liang et al., 2024; Isaacson and Scanziani, 2011), to our knowledge, none of these studies systematically explored the underlying anatomical ratios between the excitatory and inhibitory populations in recurrent neural networks.

When individual neurons are modelled as rate units, they typically do not obey Dale’s law (Eccles, 1976): their outgoing connection weights are most often drawn from a random distribution centred around zero (Sussillo and Abbott, 2009; Boström et al., 2013; Jaeger, 2001), causing them to exert both excitatory and inhibitory influences. In spiking neural networks, the neurons do obey Dale’s law, but the ratio between the number of excitatory and inhibitory neurons remains relatively unexplored. Most studies use the same 4:1 excitatory-to-inhibitory ratio to reflect the population sizes found in the cortex (Marom and Shahaf, 2002; Sahara et al., 2012; Marín, 2012; Wonders and Anderson, 2006; Meinecke and Peters, 1987). Furthermore, none of these studies focused on motor pattern generation, where spinal cord-specific EI ratios might be more relevant. Findings in rats indicate the anatomical EI ratio might be smaller in the spinal cord (Todd and Sullivan, 1990). This indicates that the 4:1 excitatory-to-inhibitory ratio is not omnipresent, and that inhibition might play a larger role in spinal motor pattern generation. This indicates that a more nuanced understanding of the excitatory-to-inhibitory balance is needed for spinal motor control.

The goal of our paper is to study the role of recurrent inhibition and global anatomical EI balance in spinal motor pattern formation. We adapt an existing reservoir computing model (Sussillo and Abbott, 2009) to have distinct excitatory and inhibitory neural populations, and introduce a new composite measure (*imbalance*) that quantifies their relative influence on the network dynamics in terms of the population’s sizes and connectivities. We initialize the networks with different anatomical imbalances and train them to reproduce muscle activation patterns calculated from human locomotion recordings. We analyze the effects of the imbalance on the network dynamics and muscle activation patterns generated by the network.

## Methods

We modelled pattern formation in the spinal cord by training simulated neural networks to produce locomotor muscle activations for 17 muscles obtained from human experiments. We investigated the influence of the ratio of excitation to inhibition in these networks by varying the numbers of neurons (*N*), connection percentages (*p*), and connection strengths (*g*) for the two populations. We relate the network performance to the intrinsic network dynamics, obtained from both our simulations and from a theoretical assessment using mean-field theory (Sompolinsky et al., 1988; Rajan et al., 2010; Mastrogiuseppe and Ostojic, 2017, 2018).

Simulations and data analysis were performed in Matlab (Version R2023a, The MathWorks, Inc., Natick, Massachusetts, United States).

### 2.1 Network

The investigated networks consisted of an excitatory population *E* with neuron number *N*_*E*_ (fig. 1, red triangles in the reservoir) and an inhibitory population *I* with neuron number *N*_*I*_ (fig. 1, blue circles in the reservoir). Any given neuron *i* in either population receives external input, output feedback and recurrent input.

**Fig. 1.**
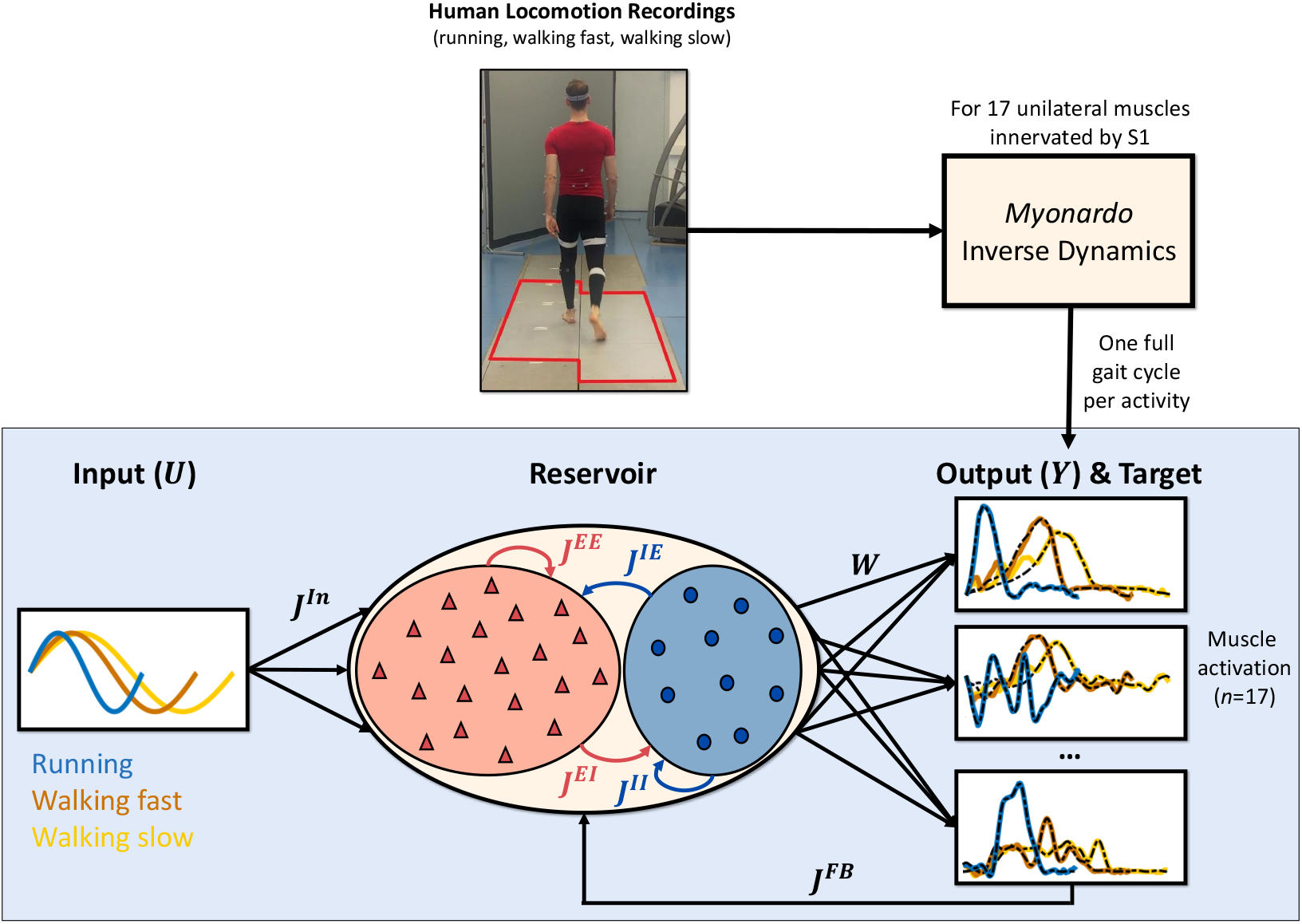
Schematic overview of the methods. Human locomotor signals were recorded from one adult during slow walking, faster walking, and running (top middle). Ground reaction forces were recorded using six Kistler Force Platforms (outlined in red) and kinematics were recorded using a Qualisys measuring system. The recorded kinetics and kinematics were then input into the musculoskeletal model *Myonardo* for inverse dynamics calculations (top right), which provided the required muscle activations for 17 unilateral muscles. These muscle activations were used as the (17-dimensional) target output of the neural network (bottom right). The neural network (bottom middle) consists of two connected excitatory (red, triangles) and inhibitory (blue, circles) populations, of which the neuron number, connection percentage and connection strength have been varied separately. The current figure shows an example network that has more excitatory than inhibitory neurons. In the network, blue lines denote inhibitory connections, while red lines denote excitatory connections. The input to the network (bottom left) was a sinusoid with a frequency corresponding to the gait frequency for each activity: running (blue), fast walking (orange), and slow walking (yellow). The muscle activation outputs resulted from all-to-all connections between the neurons in both populations via trained output weights *W*. The network outputs were fed back into the network via feedback weights *J* ^*F b*^. Example outputs are shown for all three locomotor activities (coloured lines) overlayed on the target outputs (dashed black lines) for three of the 17 output signals (from top to bottom: m. gastrocnemius, m. flexor digitorum longus, and m. gluteus medius)

The external input represents the output of the rhythm generation layer and equals *J*^*In*^*U* (*t*), where *U* (*t*) = 1 + sin(*ωt*) with *ω* equal to the target stride frequency (fig. 1, bottom left), and *J*^*In*^ the input weights drawn from a Gaussian distribution with mean 0 and variance 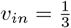. The mean and variance of the distribution were chosen to match those of Sussillo and Abbott (2009). While Sussillo & Abbott employed a uniform distribution for their input weights, we used a Gaussian distribution to align with the mean-field theory, which assumes Gaussian-distributed inputs.

The output feedback to the network equals 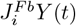, where *Y* (*t*) is the 17-dimensional output of the network (fig. 1, bottom right) and the feedback weights *J*^*Fb*^ are drawn from a uniform distribution between -1 and 1.

For the recurrent inputs within the network, we use a sparse connection matrix, reflecting the sparse connectivity of biological neural networks. The connection weights 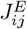 and 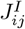 have a probability of *p*_*E*_, resp. *p*_*I*_, of being non-zero. To ensure Dale’s law, the weights for the non-zero connections are drawn from either a positive (for excitatory neurons) or negative (for inhibitory neurons) truncated Gaussian distribution, with mean zero and variance 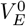. The variance 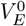 is chosen such that the variance *v*_*E*_ of the full set of excitatory weights is independent of *p*_*E*_ and scales with *N*_*E*_ according to 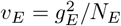 (see section A.5), where *g*_*E*_ is the parameter that scales the excitatory connection strength (Sompolinsky et al., 1988; Sussillo and Abbott, 2009). Likewise, inhibitory weights 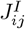 have a probability *p*_*I*_ of being non-zero. If non-zero, their value is the negative of the absolute value of a Gaussian random variable with mean zero and variance 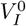, chosen such that the variance of the inhibitory weights is 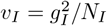.

The internal state *x*_*i*_ of each neuron *i* is initialized randomly, drawn from the standard normal distribution, and follows the dynamics:

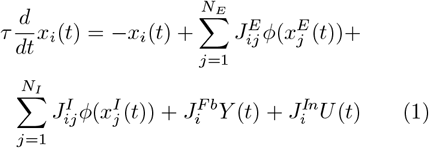

with time constant *τ* = 0.01. The neurons’ firing rate equals *ϕ*(*x*_*i*_). For biological realism, the transfer function *ϕ* should result in (1) positive firing rates, (2) zero firing rate for negative input, as biological neurons do not fire when they receive negative or zero input, and (3) saturated firing rates for large positive input, matching experimental findings of firing rate saturation (Prinz et al., 2003). Therefore, we chose the rectified hyperbolic tangent, where *ϕ* is tanh(·) if *x*_*i*_ is positive, and zero otherwise. Provided requirement (1) is satisfied, the choice of transfer function is not critical. The paper’s main results were duplicated when using a sigmoid transfer function (see fig. 11).

The output of the network is calculated from the neuron firing rates as follows:

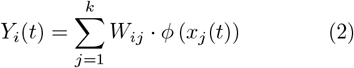

where *W*_*ij*_ is the output weights matrix, which is initialized as zeroes and evolves throughout training (see section 2.3.3 and section B).

Our network is an adaptation of an echo state neural network (Sussillo and Abbott, 2009) with two major modifications. First, synaptic weights from any given neuron are either all positive or all negative, whereas, in the original network, all recurrent weights are drawn from a Gaussian distribution of mean zero and variance 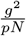, therefore a given neuron can have both positive and negative output weights (Sussillo and Abbott, 2009). Second, the firing rates of all neurons were constrained to be positive, whereas the firing rate is tanh(*x*_*i*_) in the original network, which can be either positive or negative depending on the sign of *x*_*i*_. As a result, individual neurons in our network can be considered either excitatory (red triangles in fig. 1) or inhibitory (blue circles in fig. 1).

### 2.2 Mean-field theory analysis

We performed a theoretical analysis of the networks using the mean-field theory, a technique where a description of the network dynamics can be derived self-consistently by averaging over the random parts of the network connectivity (Sompolinsky et al., 1988; Rajan et al., 2010; Mastrogiuseppe and Ostojic, 2017, 2018). For simplicity, we omitted the output feedback term in this study, but see Mastrogiuseppe and Ostojic (2018) for a full treatment of the output feedback. A detailed description of our MFT approach can be found in section A, but we will present a short overview here.

In the MFT approach, the input to each neuron is modelled as a Gaussian random process. By averaging across different realizations of the random connectivity (denoted by [.]), we can express the mean *µ* and the variance Δ of the input as a function of the mean firing rate [*ϕ*] and mean squared firing rate [*ϕ*^2^] (see Appendix section A.2) as follows:

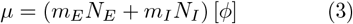

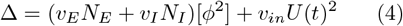

where *m*_*E*_ and *m*_*I*_ are the mean values of the excitatory and inhibitory weights, and *v*_*E*_ and *v*_*I*_ are their variances. Note that the mean-field predictions are generally valid for any variances and transfer functions, and can easily be adapted.

The population activity thus depends critically on two parameters: the mean *m*_*E*_*N*_*E*_ + *m*_*I*_ *N*_*I*_ and the variance *v*_*E*_*N*_*E*_ + *v*_*I*_ *N*_*I*_ of the sum of the recurrent weights to each neuron. We define the first parameter as the imbalance *A*:

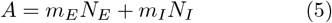

By definition, the mean weight of the excitatory connection *m*_*E*_ is positive, and the mean weight of the inhibitory connections *m*_*I*_ is negative, resulting in a total imbalance that can either be positive, in networks dominated by excitation, or negative, in networks dominated by inhibition. Our mean-field analysis investigated ranges from -15 to +15, to match the values from our simulations (described in section 2.3.1).

Next, we assumed that the network is in a quasi-stationary state, i.e., the input signal *U* (*t*) = 1 + sin (*ωt*) varies slowly compared to the time constant *τ*. Since neural integration occurs over a much faster timescale (*τ* = 10 ms) than locomotion (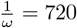 ms for running), this assumption provided a good match between theory and simulations (see fig. 16). In the quasi-stationary state, the activation *x*_*i*_ of each neuron has the same mean and variance as its input. We explicitly modelled this activation as a Gaussian variable of mean *µ* and variance Δ. In this way, we can express [*ϕ*] and [*ϕ*^2^] as a function of the neural input mean *µ* and variance Δ (see Appendix section A.3):

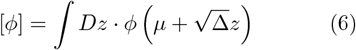

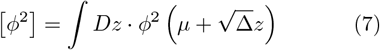

where 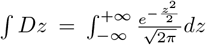 is derived from the probability distribution of the Gaussian process *z*.

We numerically solved this set of four equations with four unknowns to self-consistently determine the stationary solution given by *µ*_0_, Δ_0_, [*ϕ*]_0_, [*ϕ*^2^]_0_.

Finally, we performed a stability analysis and showed that the stationary solution loses stability for large values of *v*_*E*_*N*_*E*_ + *v*_*I*_ *N*_*I*_ (see Appendix section A.6). We determined the maximal value of *v*_*E*_*N*_*E*_ + *v*_*I*_ *N*_*I*_ for which networks are stable for all values of imbalance *A* and restricted our simulations to this stable range (i.e. *g*_*tot*_ ≤ 2.25, see fig. 12a). Note that *v*_*E*_*N*_*E*_+ *v*_*I*_ *N*_*I*_ is varied via the overall strength 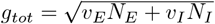, as explained in section 2.3). We find that, with this restriction, network performance does not depend on *v*_*E*_*N*_*E*_ +*v*_*I*_ *N*_*I*_, or equivalently, *g*_*tot*_ (see fig. 15a).

### 2.3 Simulations

The differential equation governing the change in neural activation (eq. (1)) was solved numerically using Euler-forward integration to get the new neuron activations for each time step (Δ*t* = 0.005*s*).

#### 2.3.1 Network parameters

For the specific weight distributions in our networks (see section A.5), the anatomical imbalance can be expressed as a function of the neuron number (*N*), the connection percentage (*p*), and the connection strengths (*g*) of the two populations according to:

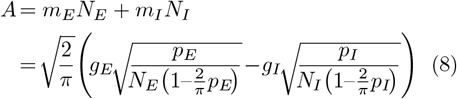

We investigated networks with imbalances ranging from − 15 to +15 in steps of 1, by globally varying either *N, p*, or *g* for both populations. This analysis was carried out for various levels of the total network size *N*_*tot*_ = *N*_*E*_ + *N*_*I*_, the mean connection percentage 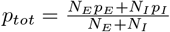, or the total connection strength 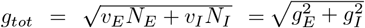.

First, the imbalance was modified via *N*_*E*_ and *N*_*I*_, with total network sizes of *N*_*tot*_ = 300, 400, 500, 750, 1000, 1500 and 2000. We chose a minimum network size of 300 as networks with fewer neurons were found to have poor performance (see fig. 15). The connection percentage was set constant at *p*_*E*_ = *p*_*I*_ = 0.1, corresponding to connection ratios found in *ex vivo* neural networks (Marom and Shahaf, 2002), and the connection strength at *g*_*E*_ = *g*_*I*_ = 1.5, as this has been shown to produce good network performance (Sussillo and Abbott, 2009).

Second, the imbalance was modified via *p*_*E*_ and *p*_*I*_, with the mean connection percentage *p*_*tot*_ ranging from 0.05 to 0.5 in steps of 0.05. The total neuron number was set constant at *N*_*tot*_ = 750, with neuron ratios varying from 4:1 to 1:4, namely: *N*_*E*_:*N*_*I*_ = 600:150, 500:250, 375:375, 250:500 and 150:600. The connection strength was set constant at *g*_*E*_ = *g*_*I*_ = 1.5.

Finally, imbalance was modified via *g*_*E*_ and *g*_*I*_, with the total connection strength *g*_*tot*_ ranging from 0.5 to 2.25 in steps of 0.25. Additionally, *g*_*tot*_ ≈ 2.12 was included, corresponding to *g*_*E*_ = *g*_*I*_ = 1.5, i.e. the connection strengths used when varying *N* and *p*. Networks with *g*_*tot*_ ≥ 2.5 were excluded from further analysis as these were predicted to be unstable by the mean-field theory analysis (see fig. 12). The total network size was set constant at *N*_*tot*_ = 750, with neuron ratios varying from 4:1 to 1:4, as above. The connection percentage was set constant at *p*_*E*_ = *p*_*I*_ = 0.1.

For each combination of parameter settings, twenty networks were trained and tested. The results were averaged over these twenty networks to decrease the influence of random initialization.

#### 2.3.2 Locomotor muscle activations

The neural networks presented in the previous sections were trained to produce locomotor muscle activations obtained from human experiments.

Kinematics and ground reaction forces were recorded for three locomotor activities performed at self-chosen speeds: slow walking, fast walking, and running. Kinetics were recorded using six Kistler force platforms (Type 9287CA, 90×60 cm, Kistler Instrumente AG, Winterthur, Switzerland – outlined in red in the photo in fig. 1). Whole-body kinematics were recorded at 200 Hz (Qualisys, Gõteborg, Sweden) using a modified Plug-In Gait marker set (Vicon Motion Systems Ltd, Oxford, UK). A single participant (male, 26 years old, 1.96m tall, weighing 86kg) walked and ran across the length of the force plates. For each of the three activities, a single stride from a right heel strike to the following right heel strike was selected. The heel strikes were detected by finding the lowest point of the right heel marker, as recorded by the Qualisys system. This data was obtained as part of another experiment that has been approved by the local Ethics Committee of the Faculty of Psychology and Sports Science at the University of Münster (#2019-10-RD). The participant signed the informed consent form before the start of the measurements.

Muscle activations, ranging from 0 to a theoretical maximum of 1, were calculated from the recorded kinetics and kinematics using the inverse dynamics function of the musculoskeletal model *Myonardo®* (Predimo GmbH, Münster, Germany). For more information on the musculoskeletal model and the performed calculations, see section C and Wagner et al. (2022). From the model output, we selected the 17 muscles on the right side of the body that are innervated by the S1 spinal segment (Sharrard, 1964): m. gluteus maximus, m. gluteus medius, m. gluteus minimus, m. biceps femoris (caput longum and breve), m. semitendinosus, m. semimembranosus, m. tensor fasciae latae, m. piriformis, m. gastrocnemius, m. soleus, m. peroneus longus and brevis, m. flexor hallucis longus, m. flexor digitorum longus, m. extensor hallucis longus, and m. extensor digitorum longus. This resulted in a 17-dimensional target signal for each of the 3 locomotor activities.

These single-stride target signals were then used as building blocks to assemble the full target signals used to train and test the networks. The training target signal consisted of 15 strides, with each of the three activities repeated for five consecutive strides. The test target signal consisted of 21 strides, with the three activities randomly interleaved. A different test signal was created for each of the twenty network instances, by changing the sequence. The same set of twenty signals was used across all parameter settings. The transitions between two consecutive strides were smoothed by leaving a one-sample gap between the two strides, which was then filled through linear interpolation. The full target signals were then filtered using a 20 Hz low-pass bi-directional Butterworth filter (2^nd^ order). Finally, the first and last 50 samples were cut from the full signals to remove any unwanted filter artifacts. Only the first 20 cycles were used to evaluate network performance, thereby excluding the transient phase of the network from the evaluation.

#### 2.3.3 Training and testing the networks

The output weights of the networks (*W*_*ij*_) were trained using the recursive least squares algorithm (Haykin, 2014, see also section B). Unlike the internal connection weights, the output weights of the network were not restricted to be positive during training and can be either negative or positive. All networks were trained over 5 consecutive training iterations, with the network activations and their derivatives reinitialized at the start of each iteration.

#### 2.3.4 Outcome parameters

##### Network Dynamics

Network dynamics were assessed by recording the mean and neuron-to-neuron (or network) variance of the recurrent input *x* and the firing rates *φ*. This network variance provides a measure for the firing rate heterogeneity, reflecting the diversity of neuronal activity across the population.

##### Performance quantification

The performance of each network was quantified as the percentage of successfully reproduced strides. First, the root-mean-square error (RMSE) between the actual (coloured lines, fig. 2) and target (black lines, fig. 2) output was calculated for each muscle and each of the 20 strides in the test signal. The pattern produced by each muscle during each stride was then classified as either a success if RMSE ≤ 0.05 (shaded in green in fig. 2) or a failure if RMSE *>* 0.05 (shaded in red in fig. 2).

**Fig. 2.**
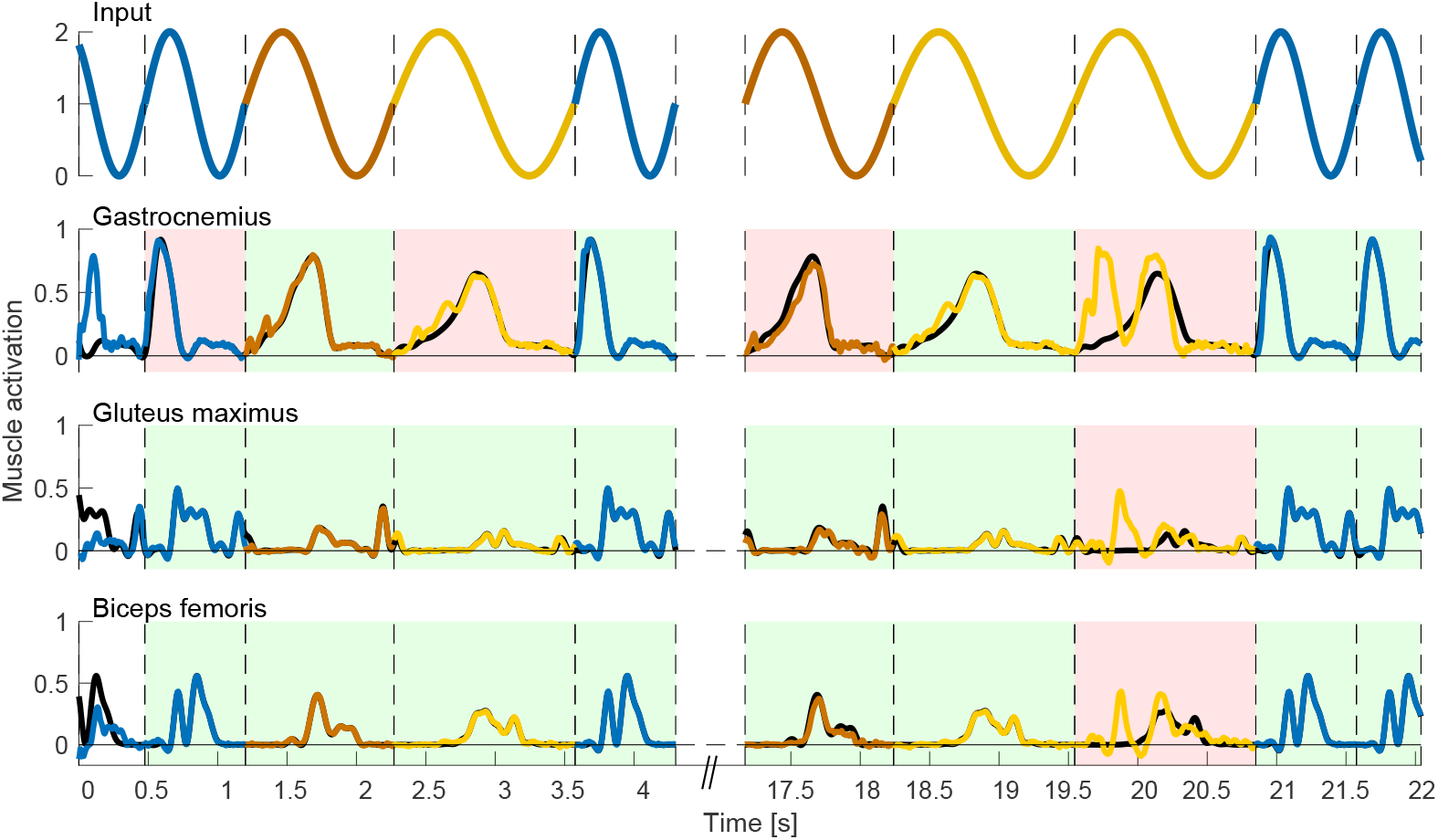
Performance calculation in example output signals. The networks are tested on their ability to reproduce the motor patterns of randomly interleaved gait cycles. The input to the network (top row) and network outputs of three example muscles (bottom rows) are shown for slow walking (yellow), fast walking (orange) and running (blue). The target output is indicated by the solid black line (bottom rows). The transitions between cycles are indicated as dashed vertical black lines. For each cycle and muscle, the cycle is classified as a success (green background) if the root mean square difference between the target and the actual output is less than 0.05. Otherwise, it is classified as a failure (red background). The performance of the network is the percentage of successful cycles over the full set of 17 muscles and 20 cycles. Note that the first cycle is not taken into account for the performance evaluation (white background) as it covers the initialization period of the network. This figure shows the output of a random network with *N*_*E*_ = *N*_*I*_ = 375, *p*_*E*_ = *p*_*I*_ = 0.1 and *g*_*E*_ = *g*_*I*_ = 1.5 and a performance of 78.2% (66.7% if only taking into account the visible cycles)

##### Effective network dimensionality

We quantified the effective dimensionality of the internal network dynamics as the number of network principal components (NPCs) necessary to account for 99% of the variance in the neuron firing rates. The effective network dimensionality characterizes the complexity of the network dynamics by indicating the number of unique components that are needed to capture the majority of the variance in its activity. Low dimensionality indicates that the neuronal activity patterns in the network are highly correlated with each other (as in the right-hand side panels of fig. 17), whereas high dimensionality indicates diverse neuronal activity, which allows the network to capture more complex patterns (as in the left-hand side panels of fig. 17). The same approach was applied to quantify the number of principal components in the output and target output signals.

##### Muscle parameters

We calculated the coactivation index (*CAI*) between the muscle activation output patterns. For every muscle pair, the coactivation can be calculated as (Falconer, 1985; Souissi et al., 2017):

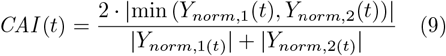

Here, *Y*_*norm*,1_ and *Y*_*norm*,2_ are the activations of both muscles, normalized to their own maximum over the evaluated time period. The coactivation index is reported as an average over the entire test signal.

##### Output Variance

We additionally calculated the variance between the different locomotor activities, to assess the network’s ability to generate distinct signals for each task. After time-normalizing all strides from 0 to 100%, we computed the mean and variance across cycles for each of the three tasks individually. We could then determine the between-task variance by calculating the variance over the three resulting averaged cycles.

## 3 Results

We present our findings on the impact of the global anatomical excitatory to inhibitory imbalance in recurrent neural networks trained to replicate locomotor signals. Our results include a theoretical assessment of network dynamics using mean-field theory, complemented by simulation analyses that explore the relationship between the imbalance, effective dimensionality, and motor outputs.

### 3.1 Mean-field theory

The mean-field predictions for the mean and variance of both the recurrent input and the firing rate showed a good match to simulations for all studied parameters (see figs. 3 and 16). The MFT analysis showed that, for the analysed parameter ranges, the internal network dynamics do not depend on the total number of neurons or the mean connection percentage, but only on the imbalance (*A*) and the total connection strength (section A, eqs. (38) and (39)). While the total connection strengths have quantitative differences, they do exhibit qualitatively similar behaviour (fig. 16).

**Fig. 3.**
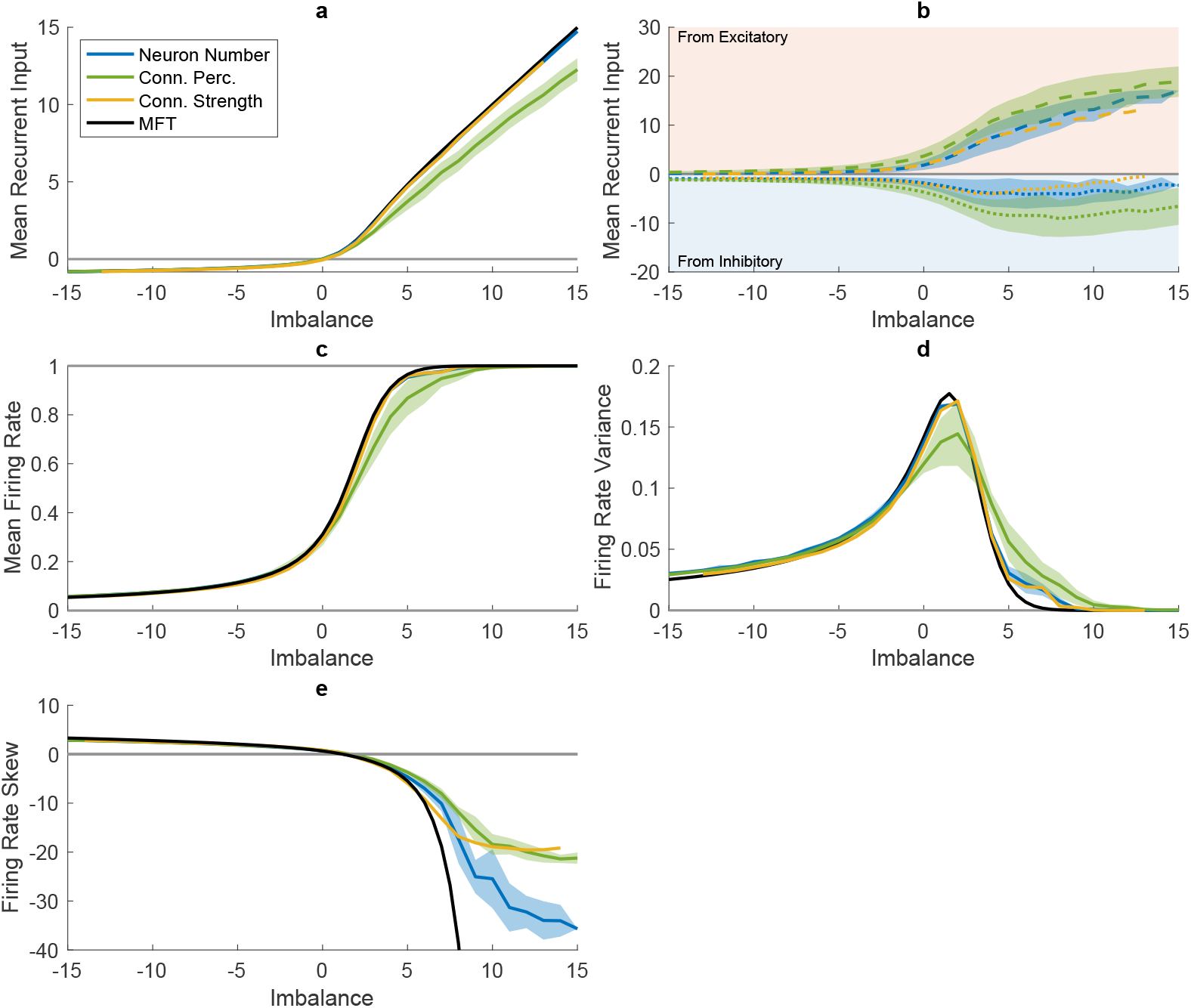
Network Dynamics as a function of imbalance A, with (a) showing the mean of the input to individual neurons (*x*), (b) the mean input split between the input coming *from* the excitatory (positive; in the red-shaded area) and inhibitory (negative; in the blue-shaded area) populations, (c) the mean of the firing rate (*φ*), (d) the firing rate variance, and (E) the skew of the firing rate distribution. The averages over all levels of *N*_*tot*_ (blue) and *p*_*tot*_ (green) are depicted, with the shaded area showing the standard deviation. Only one level of *g*_*tot*_ is shown (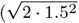, matching the total connection strength used when varying the neuron number and connection percentage) as the total level influences the relationship between imbalance and the firing rate (see fig. 16). The black lines show the mean-field theory (MFT) predictions for each imbalance with 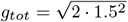. All simulation results have been averaged over 20 network instances

### 3.2 Network Dynamics

As predicted by our mean-field theory analysis (fig. 3, black), we found that the network dynamics depend critically on the excitatory to inhibitory imbalance, regardless of whether it was varied in terms of neuron number (fig. 3, blue), connection percentage (fig. 3, green) or connection strength (fig. 3, yellow). The effect of the imbalance is asymmetrical: increasing the inhibition has a markedly different effect than increasing excitation.

When increasing the excitation in the network, we see a rapid, almost linear increase of the recurrent input with imbalance *A* (fig. 3a). Here, the incoming excitatory current (dashed lines in fig. 3b) overpowers the incoming inhibitory current (dotted lines in fig. 3b). This large excitatory current causes the mean firing rate to saturate (fig. 3c), which not only keeps the neuronal input high, but also results in a drastic drop in the firing rate variance (fig. 3d, with most or all neurons firing at their maximum constantly (see fig. 17, *A* = 5, 7.5 and 10). This is also evidenced by a strong negative skewness in the firing rate distributions (see fig. 3e, and example histograms in fig. 18).

In contrast, for inhibitory networks (*A <* 0), the mean input received by the neurons is negative but small (fig. 3a), with the recurrent inputs from both the excitatory and inhibitory populations relatively balanced and close to zero (fig. 3b). As a results, small variations in the inputs to individual neurons can drive variations in the neurons’ firing rate, thus preventing the firing rates from saturating at their minimum value of zero (fig. 3c and fig. 17, *A* = − 5 and 0). Consequently, the firing rate variance does not drop to zero (fig. 3d) and the firing rate distribution has a moderately positive skew (fig. 3e).

### 3.3 Dimensionality and Performance

Varying the imbalance led to a large variety in the effective dimensionality of the networks, with the number of NPCs ranging from 1 to 171 in individual networks. Due to the low firing rate heterogeneity in excitation-dominated networks, the dimensionality of the network drops drastically when excitation increases (fig. 4a and fig. 17, *A* = 5, 7.5 and 10). This is accompanied by a concurrent drop in network performance (fig. 4b). While the dimensionality in inhibitory networks is also lower than in balanced networks (fig. 4a), the number of principal components in the network stays well above the number of principal components in the target data (i.e. 9 PCs explaining *>* 99% of the output variance, as indicated by the pink line in fig. 4a,c). This means that the effective dimensionality of inhibition-dominated networks is theoretically large enough to reproduce all dimensions present in the target output, as reflected by their relatively high performance (fig. 4b).

**Fig. 4.**
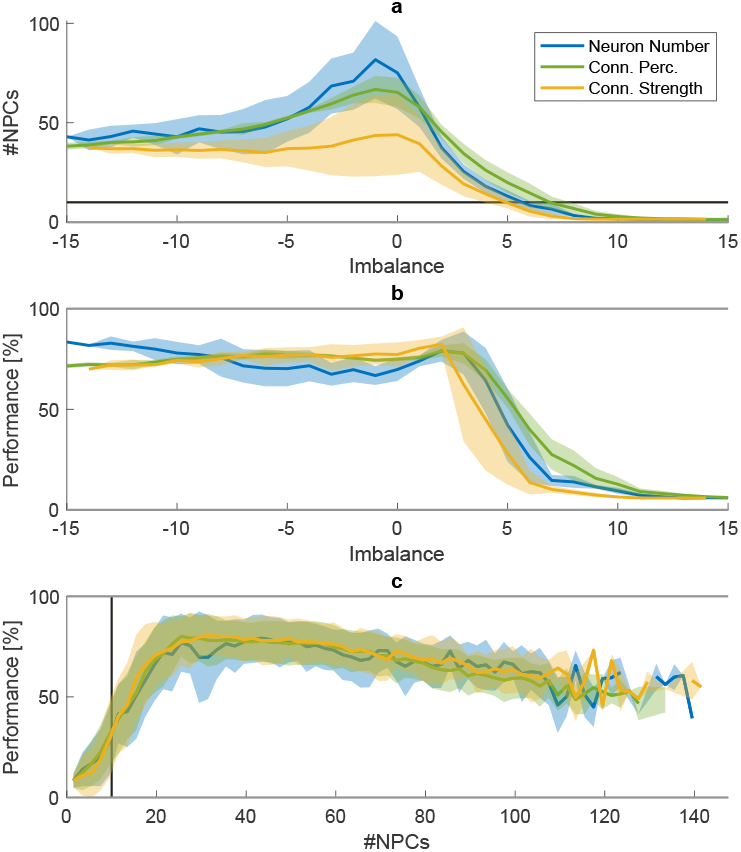
Effective dimensionality and performance of the networks. The imbalance *A* is plotted against (a) the number of network principal components (#NPCs) and (b) the network performance. (c) depicts the network performance as a function of the number of NPCs. The average over 20 network instances and all included levels of *N*_*tot*_, *p*_*tot*_ and *g*_*tot*_ are depicted, with the shaded area showing the standard deviation over these levels. In A and C, the pink line indicates the number of principal components in the target output (i.e., 9)

Indeed, network performance depends critically on the effective dimensionality of the networks (fig. 4c). For our selection of stable networks, we see an approximately linear increase in performance as the number of NPCs grows before plateauing and slowly dropping (fig. 4c). Thus, there seems to be an optimal window for the number of NPCs around approximately 40. This corresponds to 2.8-6.4 times the number of output principal components. Although the influence of the imbalance on the effective dimensionality depends on *g*_*tot*_, *N*_*tot*_, and *p*_*tot*_ (see fig. 19), the influence of the imbalance on performance is largely independent of these factors (fig. 15, solid lines). This difference can be explained by the relatively wide plateau and slow drop-off in fig. 4c), indicating that, above a certain threshold, a change in the number of effective dimensions does not necessarily lead to a large change in performance.

### 3.4 Motor patterns

Shifting the imbalance toward inhibition also produces a markedly different effect on the motor output compared to shifting it towards excitation. fig. 5 shows the target as well as the actual network motor outputs for four example muscles for slow walking (red), fast walking (yellow), and running (blue). Balanced networks and networks with negative imbalance (first and second column in fig. 5) can produce distinct motor outputs for the three tasks. In contrast, in excitation-dominated networks (third, fourth and fifth column in fig. 5), the motor outputs for the three tasks become increasingly similar, which is also evidenced by the decrease in the variance between the locomotor tasks (see fig. 6b). In addition, the outputs of the different muscles become more similar. This is evidenced by an increase in the coactivation index *CAI* (see fig. 6a). The reduced firing rate heterogeneity and effective dimensionality in the excitatory networks are thus accompanied by a simplification of the networks’ outputs.

**Fig. 5.**
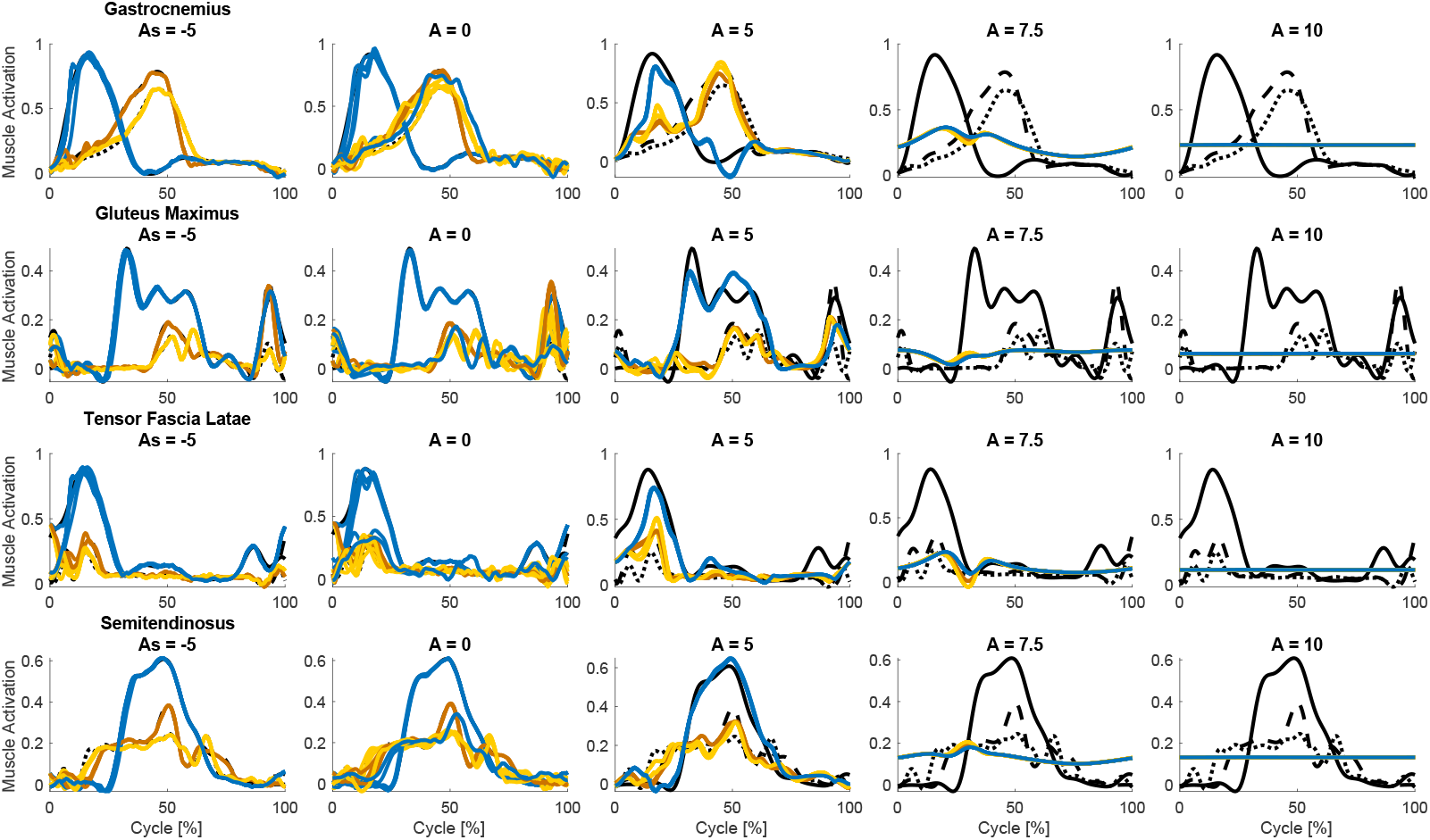
Example output signals plotted as a function of the cycle percentage for various imbalances, with 0% and 100% corresponding to right heel strikes. Each column shows signals belonging to a random example network with an imbalance of *A* = *−*5, 0, +5, and +10 from left to right. The imbalances were varied by changing the connection strengths for the two populations (from *g*_*E*_ *≈* 0.92 and *g*_*I*_ *≈* 1.91 for *A* = *−*5, to *g*_*E*_ *≈* 2.12 and *g*_*I*_ *≈* 0.14 for *A* = 10), while keeping the other parameters constant (*N*_*E*_ = *N*_*I*_ = 375, *p*_*E*_ = *p*_*I*_ = 0.1,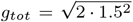). Each row shows the output corresponding to a different muscle. From top to bottom, these are the m. gastrocnemius, m. gluteus maximus, m. tensor fascia latae and m. semintendinosus. Black lines indicate the target output for each activity, with solid lines representing running, dashed lines fast walking, and dotted lines slow walking. Coloured lines indicate the actual network output, where blue depicts running, yellow fast walking, and red slow walking. These example networks had a success percentage of 85%, 71%, 50%, 8% and 6% respectively

**Fig. 6.**
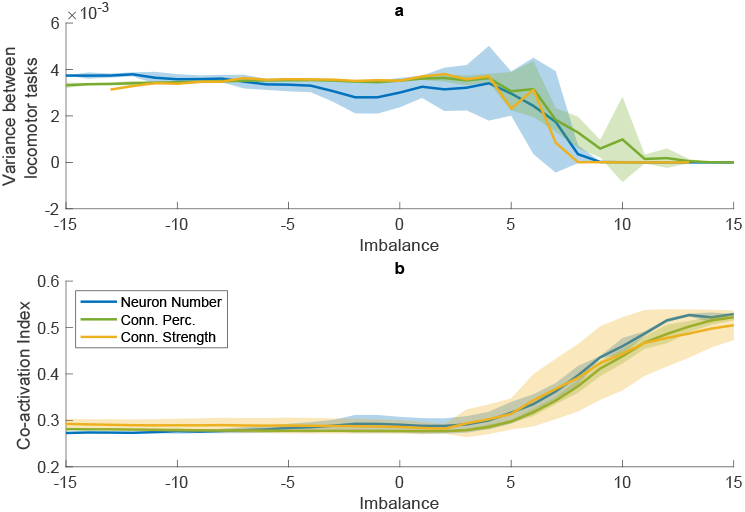
Increased excitation leads to a simplification of the motor output, both between muscles as well as between locomotor tasks. (a) average coactivation index between each pair of muscles and (b) variance between the outputs for the different locomotor tasks (i.e., slow walking, fast walking, and running), as a function of the imbalance *A*, when changing the number of neurons (blue), connection percentage (green), and connection strength (yellow). Both panels represent averages over 20 network instances and all levels of *N*_*tot*_, *p*_*tot*_, and *g*_*tot*_, with the shaded area depicting the stan<u>dard d</u>eviation over these levels. For panel A, only 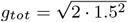 is shown, as the overall connection strength influenced the relationship between the imbalance and the variance (see fig. 16c)

## 4 Discussion

This study aimed to explore how the global ratio of recurrent inhibition to excitation affects the dynamics, performance and motor outputs of RNN-based CPG models, with a focus on pattern formation. Reservoir computing models were trained to recreate muscle activation patterns derived from human locomotion, using various ratios of recurrent excitation to inhibition. We performed a mean-field analysis and derived a composite measure called imbalance that quantifies the overall ratio of inhibition to excitation of the recurrent connections in the network. The mean-field analysis predicted that a high imbalance (i.e. where excitation ≫ inhibition) would lead to saturated firing rates and, thus, to reduced network variance. Conversely, inhibition-dominated networks were expected to maintain higher network variance. Our simulations confirmed these predictions and additionally showed that inhibition is necessary both for good network performance and to recreate experimentally observed firing rate distributions. In contrast, networks with increased excitation tended to produce simplistic and repetitive motor patterns.

Many studies have investigated the balance between excitation and inhibition, both in biological and artificial neural networks (reviewed in, e.g., Hennequin et al., 2017; Sadeh and Clopath, 2021; Herstel and Wierenga, 2021; Liang et al., 2024; Isaacson and Scanziani, 2011), demonstrating that a functional EI balance is crucial for proper neural functioning. However, this functional balance does not necessarily result from a strict underlying anatomical balance or symmetry. In biological networks, the anatomical characteristics of excitatory and inhibitory neurons do not follow a 1-to-1 ratio. For instance, there are typically more excitatory than inhibitory neurons: inhibitory neurons make up only 30 to 45% of the neurons in laminae I to III of the rat spinal cord (Todd and Sullivan, 1990), and only around 20% of the neurons in the mammalian cerebral cortex (Marom and Shahaf, 2002; Sahara et al., 2012; Marín, 2012; Wonders and Anderson, 2006; Meinecke and Peters, 1987). The connection probabilities are also not equal, as, in the cortex, only one in fifteen inputs to a given neuron is inhibitory (Megías et al., 2001; Peters, 2002). Conversely, inhibitory neurons have been found to synapse closer to the axon somata (Beaulieu et al., 1992; Peters, 2002), thereby increasing their effectiveness relative to the excitatory neurons (Chen et al., 2012; Markram et al., 2004). It is unclear, however, whether this compensates for the smaller number of connections, or whether there still is a resultant anatomical imbalance between excitation and inhibition. In this study, we examined the influence of the anatomical ratio between excitatory and inhibitory recurrent connections and showed that, as a natural consequence of the recurrent input, the net current to the neurons can still be balanced, even if the individual anatomical parameters are asymmetrical or imbalanced.

To investigate the effects of the anatomical EI balance, we introduced the anatomical *imbalance* to quantify the excitatory and inhibitory populations’ influence in one single variable. The imbalance has the same impact on the network dynamics, regardless of which of the underlying population characteristics (neuron number, connection percentage, and connection strength) are adjusted. The described relationship between the imbalance, network dynamics and performance only breaks down when other factors disrupt network functioning, e.g., when the connection strength rises beyond the stable region (see fig. 15a, *g*_*tot*_ ≥ 2.5), or when the networks become too small (fig. 15b, *N*_*tot*_ ≤ 250). The imbalance and strength equations are expressed in terms of the mean, variance, and size of the two neuron pools. Therefore, the equations can readily be adapted to networks with different distributions and scaling factors, and we expect our results to generalize to these networks.

Our findings indicate that recurrent inhibition is not only important to maintain network performance, but also vital in replicating features observed in experimental recordings. First, recurrent inhibition is necessary to recreate the positively skewed firing rate distributions observed in experimental studies. Specifically, networks must exhibit a slight anatomical imbalance favouring inhibition to recreate experimentally observed firing rate skewness. Excitation-dominated networks have negatively skewed distributions, with a prominent peak at the maximum firing rate. Increasing the inhibition broadens the distributions and causes the peak to shift leftward, thereby matching the lognormal firing rate distributions that have been experimentally recorded in the spinal cord (Petersen and Berg, 2016) and various cortical areas (Shafi et al., 2007; Hromádka et al., 2008; O’Connor et al., 2010) across different species. These findings are consistent with previous works, both computational and experimental, that showed that balanced networks have firing rate distributions that are relatively wide and skewed at low average firing rates (Hennequin et al., 2017; Vogels et al., 2005; van Vreeswijk and Sompolinsky, 1996; Lindén and Berg, 2021; Roxin et al., 2011), whereas excitatory networks exhibit a distribution with a single peak at the maximum firing rate (Lindén and Berg, 2021). Similarly, Roxin et al. (2011) showed *in silico* that increasing the inhibitory neurons’ firing rate via the external input transforms the firing rate distribution from negatively skewed to a more lognormal distribution, thereby matching experimental results recorded *in vivo*. This shift means that balanced networks can respond to external inputs in a graded manner, rather than the all-or-none firing rate seen in excitatory networks (Lindén and Berg, 2021). Our study adds that the same is true for inhibition-dominated networks.

Second, inhibition is important to prevent a loss of the effective dimensionality of the network. The network dimensionality is crucial for network performance (Susman et al., 2021; Carroll and Pecora, 2019; Carroll, 2020). In our study, networks with too few principal components could not replicate the target output signals. This matches prior findings, where increasing the rank of the reservoir matrix (which places an upper bound on the number of meaningful principal components that can be extracted) is associated with a decrease in the test error (Carroll and Pecora, 2019). In excitatory networks, the effective reservoir dimensionality is reduced below the target output dimensionality and insufficient to reproduce the required output patterns. This is caused by run-away activity in the network leading to saturated firing rates, which can be stabilized by sufficient inhibition (Sadeh and Clopath, 2021; Herstel and Wierenga, 2021). These results highlight the importance of recurrent inhibition in maintaining appropriate effective network dimensionality.

Third, our results indicate that inhibition plays a crucial role in generating diverse and variable motor patterns. High excitation networks could not produce different outputs for different muscles, as evidenced by the high coactivation and low number of output principal components. Nor were they able to differentiate between the different locomotor tasks, as evidenced by the drop in the between-task variance. These findings align with *in vivo* studies, where blocking inhibition led to synchronous activation of flexor and extensor motoneurons (Beato and Nistri, 1999; Cazalets et al., 1998; Cowley and Schmidt, 1995; Kiehn, 2006).

The importance of inhibition in the generation of diverse movements is also supported by the reduced complexity of muscle activation patterns observed in various motor disorders. In cerebral palsy, there is evidence of a decreased inhibition (reviewed by Fogarty, 2023) as well as a reduction in the number of motor modules during walking (Steele et al., 2015; Tang et al., 2015; Bekius et al., 2020), indicating a reduced locomotor complexity. Condliffe et al. (2016) showed that poor motor ability in cerebral palsy is associated with reduced inhibitory control of motoneurons. Similarly, stroke patients also show a loss of motor modules (Clark et al., 2010; Cheung et al., 2012), which is associated with reduced walking speed (Routson et al., 2013) and a decreased ability to adapt speed, timing, and step characteristics in mobility tasks (Routson et al., 2014). In stroke, however, the role of inhibition is less clear, with inhibition seemingly having a protective impact post-stroke by counteracting excitotoxicity and the associated increase in EI balance (Sydserff et al., 1995), but hampering post-stroke recovery (Grigoras and Stagg, 2021). The early neuroprotective effect of inhibition is associated with improved functional ability (Marshall et al., 1999), which matches our results.

Based on these findings, we suggest that dystonia may be the result of reduced inhibition in spinal circuits causing excessive co-contraction. Dystonias are movement disorders characterized by spontaneous muscle activity at rest, large co-contraction of flexors and extensors during voluntary movements such as walking (Rothwell et al., 1983; Cohen and Hallett, 1988), and reduced reciprocal inhibition (Nakashima et al., 1989; Panizza et al., 1990). Dystonias are thought to result from basal ganglia dysfunction, but neurophysiological evidence for this has remained elusive. The most prevalent genetic form of dystonia is early-onset generalized torsion dystonia, commonly caused by a mutation of the TOR1A gene (Ozelius et al., 1999). When this gene is deleted in spinal circuits in mice, the mice develop dystonia with symptoms corresponding to those of humans (Pocratsky et al., 2023). These recent experimental results thus suggest that dystonia may be a disorder of spinal circuits, rather than basal ganglia circuits as commonly assumed. We suggest that this disorder may be due to reduced inhibition in spinal circuits causing excessive cocontraction. Indeed, the most effective treatment for dystonia is long-term deep brain stimulation, which has been shown to improve reciprocal inhibition and reduce agonist-antagonist co-contraction (Tisch et al., 2006).

Such findings are not limited to motor disorders alone. In higher neural centres, an experimental disruption of the excitatory to inhibitory (EI) balance towards excitation can result in a variety of pathologies, such as epilepsy (Dichter and Ayala, 1987; Dudek and Sutula, 2007), autism (Casanova et al., 2003; Rubenstein, 2010; Rubenstein and Merzenich, 2003; Markicevic et al., 2020) and schizophrenia (Yizhar et al., 2011; Murray et al., 2014). Similarly, disinhibition in the spinal cord has been shown to lead to allodynia (Lee et al., 2019; Yaksh, 1989). While these disorders are not classified as motor disorders, they are often accompanied by changes in motor control. In autism spectrum disorder, for example, individuals often display atypical motor behaviours, including atypical posture and gait, hypotonia, and impaired bilateral coordination, balance and fine motor skills (Zampella et al., 2021; Jansiewicz et al., 2006; de Jong et al., 2011). In schizophrenia, motor abnormalities such as catatonia have been linked to EI imbalance (Walther and Strik, 2012). While this study provides valuable insights into the importance of recurrent inhibition, several limitations must be acknowledged. First, this study was limited to three locomotor activities, measured in a single participant. Due to this limited activity range, the networks were only trained on three output frequencies. As the frequency for which the network performs optimally depends on factors such as the connection strength or the target amplitude (Susman et al., 2021), we do not know how well our results generalize to untested frequencies. It has, however, been shown that reservoir computing networks are capable of ‘morphing’ to untrained signals (Bostrõm et al., 2013). This is also the case for our network (see fig. 20), although we could not quantify its morphing ability as we did not have target data for the intermediate frequencies. Therefore, we performed an additional experiment in which we trained networks with various imbalances to recreate motor primitives, a set of temporal basis functions, and generalize them to various frequencies. Previous experimental work in humans has shown that muscle activities over a range of locomotor speeds can be accounted for by five such primitives (schematically represented in fig. 7b), each peaking at a different part of the locomotor cycle. Confirming our main results, balanced and inhibition-dominated networks were able to generalize to the untrained frequencies, while excitation-dominated networks could not (see fig. 7). When used as inputs in neuromechanical models such as those of Aoi et al. (2019) or Di Russo et al. (2023), these primitives could thus generate locomotion over a wide range of speeds. This finding underscores our network’s potential to create locomotor patterns for unseen speeds.

**Fig. 7.**
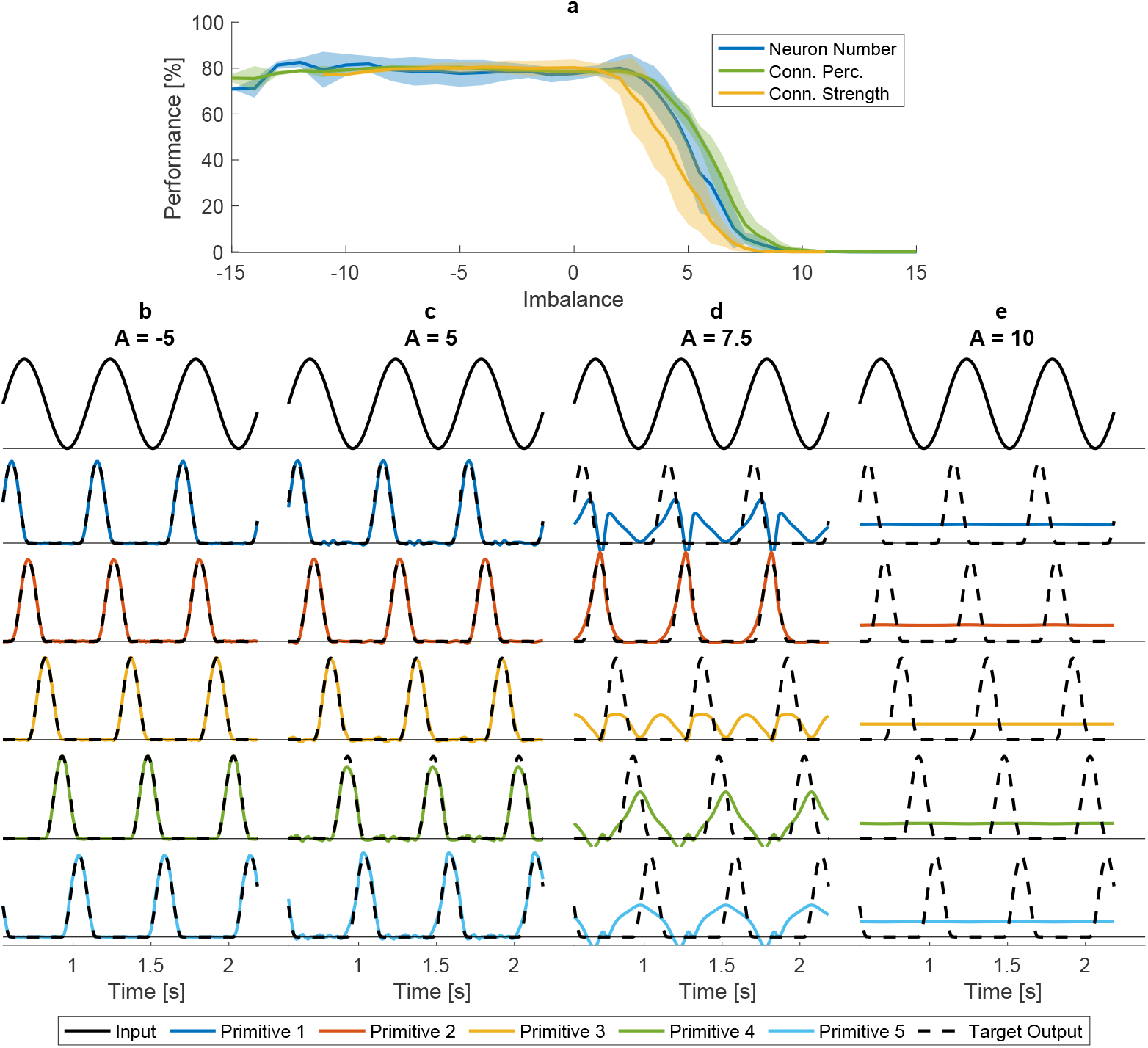
Performance on a generalization task. Networks with various imbalances *A* and total connections strengths *g*_*tot*_ were trained to reproduce 5 bell-shaped patterns representing motor primitives (based on Di Russo et al., 2023). Networks were trained on patterns corresponding to three speeds (0.6, 1.2, and 1.6 m/s) and tested on speeds ranging from 0.3 to 2.0 m/s in steps of 0.1 m/s. (a) Performance of the networks on untrained speeds, as a function of imbalance. The average over 20 network instances and all included levels of *N*_*tot*_, *p*_*tot*_, and *g*_*tot*_ are depicted, with the shaded area showing the standard deviation over these levels. (b-e) Inputs (top row), outputs (coloured lines) and target outputs (dashed black lines) for the five motor primitive patterns for networks with *N*_*E*_ = *N*_*I*_ = 375, *p*_*E*_ = *p*_*I*_ = 0.1, and 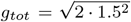 and imbalances of (b) *A* = *−*5, (c) *A* = 5, (d) *A* = 7.5, (e) *A* = 10 that were varied via *g*_*E*_ and *g*_*I*_

Second, sensory feedback, which is known to contribute to stability, perturbation responses, and even steady-state muscle activation during locomotion (Frigon et al., 2021), was not included in our model. Nonetheless, we argue that our findings most likely generalize, as feed-forward pattern formation plays a crucial role in shaping locomotor output, regardless of sensory feedback. Experimental work in deafferented animals has shown that patterns more complex than simple flexion-extension alteration could be produced in these animals without sensory input, albeit not as robustly (Grillner and Zangger, 1984; Taub et al., 1975; Grillner et al., 1995). Moreover, studies have employed neuromechanical models to explore the roles of both sensory feedback and pattern formation. These works have shown that neuromechanical models cannot rely on sensory feedback alone: a pattern formation layer in some form was needed to generate stable locomotion over a range of speeds (Geyer and Herr, 2010; Di Russo et al., 2023). Our findings demonstrate that excessive excitation impairs this intrinsic pattern formation (fig. 4b). According to the aforementioned modelling studies, adding sensory feedback is not sufficient to restore locomotion in the excitation-dominated networks. Additionally, our generalization experiment (described in the paragraph above, and in fig. 7) shows that our networks can produce the temporal primitives that are able to generate stable locomotion over a range of speeds, when used as input to a neuromechanical model with sensory feedback. We, therefore, expect our findings to generalise to a model with sensory feedback: balanced and inhibition-dominated networks will be able to produce stable locomotion, whereas excitation-dominated networks will not.

Third, our results are based on firing rate, rather than spiking, neural networks. It is unclear whether spiking networks exhibit the same robustness to inhibition as our firing rate networks, or if excessive inhibition in spiking networks would result in quiescence.

Fourth, the fixed connectivity within the neural reservoir limits the biological realism of our model. Although computationally efficient, the static reservoir connections preclude our network from synaptic plasticity, which plays an important role in maintaining functional EI balance (Vogels et al., 2011; Froemke, 2015; Field et al., 2020). This thus limits the model’s ability to simulate the dynamic adjustments observed in biological neural networks. However, for our study, this fixed structure was necessary to study the effects of the excitation-to-inhibition ratio, as neural plasticity would change the networks’ imbalance and thus skew the results. Future work could focus on the interplay between initial imbalance and synaptic plasticity.

Finally, our networks had a randomly initialized connectivity, that does not necessarily reflect the structure of the spinal neural circuits. Structured connections are often modelled in neural circuit models, which are low-dimensional and focus on the properties of specific neural subgroups. However, these models typically fail to capture the heterogeneity present within the subgroups they model. The higher-dimensional neural network models that we employ can account for this heterogeneity, but the complexity of the mathematical analysis grows with the number of subpopulations that are included. Studying the role of specific connectivities in high-dimensional network models is therefore challenging, and currently the subject of active research. Shao and Ostojic, for example, investigated the effect of various neural motifs on network dynamics and showed that adding reciprocal motifs (i.e., modeling the effect of renshaw cells) only has a small effect on network dynamics (Shao and Ostojic, 2023), whereas chain motifs have a much larger effect: increasing the strength of chain motifs can induce instability even in inhibition-dominated networks (Shao et al., 2024). However, the presence of chain motifs in the spinal cord has not been documented.

In conclusion, our study highlights the role of recurrent inhibition and anatomical EI balance in generating complex and coordinated motor patterns. Networks that were dominated by inhibition displayed robust and complex firing patterns, and were thus able to learn and reproduce human locomotor patterns effectively. In contrast, networks that were dominated by excitation exhibited a reduced effective dimensionality and simplified motor outputs. These findings demonstrate the potential for neural networks to tolerate excessive inhibition, but not increased excitation. The inclusion of sufficient recurrent inhibition was not only essential in generating the locomotor patterns; it was also necessary to reproduce experimental observations of skewed firing rates. Future work should explore a wider range of motor tasks to test and expand our findings, and investigate if they generalize to spiking neural networks.

## Supplementary information

Not applicable.

## Acknowledgements

We thank Predimo GmbH for providing the Myonardo software.

## Declarations

### Funding

This work was supported by the *Interdisciplinary teaching programme on Machine Learning and Artificial intelligence* (InterKI), via the Federal ministry of Education and Research (BMBF). Grant number: 16DHBKI049

### Competing interests

H. Wagner is shareholder in Predimo GmbH. The other authors declare that they have no competing interests that could have influenced the results of this paper.

### Ethics approval and consent to participate

The study has been approved by the Ethics Committee of the Faculty of Psychology and Sports Science at the University of Münster (#2019-10-RD). The participant signed the informed consent form before the start of the measurements.

### Consent for publication

The authors affirm that human research participants provided informed consent for publication of the images in fig. 1.

### Data and code availability

Data and code are made available at https://doi.org/10.5281/zenodo.13981334

### Materials availability

Not applicable.

### Author contribution

Conceptualization: Myriam de Graaf, Heiko Wagner; Methodology: Myriam de Graaf, Heiko Wagner, Luis Mochizuki, Charlotte Le Mouel; Formal analysis and investigation: Myriam de Graaf, Charlotte Le Mouel; Writing -original draft preparation: Myriam de Graaf, Charlotte Le Mouel; Writing - review and editing: Myriam de Graaf, Heiko Wagner, Luis Mochizuki, Charlotte Le Mouel; Funding acquisition: Heiko Wagner; Supervision: Heiko Wagner, Luis Mochizuki, Charlotte Le Mouel.

## Appendix A Mean-Field Theory

We studied the networks analytically through the application of mean-field theory (MFT). With this technique, a description of the network dynamics can be derived self-consistently by averaging over the random parts of the network connectivity (Sompolinsky et al., 1988; Rajan et al., 2010; Mastrogiuseppe and Ostojic, 2017, 2018). Following this approach, we decompose the input received by each neuron into a static component and a component that depends on the random instantiation of the network connectivity. We then average across different realisations of the network connectivity to determine the mean and variance of this random component. This allows us to self-consistently determine the mean and variance of the neuron firing rates.

### A.1 Network Dynamics

We consider a recurrent network of *N*_*E*_ excitatory and *N*_*I*_ inhibitory neurons. Each neuron *i* is characterized by its activation *x*_*i*_(*t*) and its activity *ϕ*(*x*_*i*_(*t*)), corresponding to its firing rate. The dynamics follow:

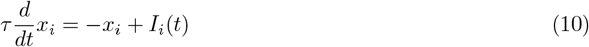

where *I*_*i*_(*t*) is the input received by any neuron *i*. This input is the sum of three terms: the external input 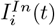, the excitatory recurrent input 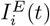, and the inhibitory recurrent input 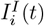. Note that we consider the network dynamics in the absence of output feedback. A derivation of the mean-field theory in the presence of output feedback can be found in Mastrogiuseppe and Ostojic (2018).

The external drive *U* (*t*) is fed to the network via input weights 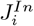, which are drawn from a random distribution with mean *m*_*In*_ and variance *v*_*In*_, such that each neuron receives external input 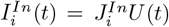. The outgoing weights from the excitatory (resp. inhibitory) neurons 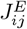 (resp. 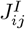) are drawn from a random distribution with mean *m*_*E*_ (resp. *m*_*I*_) and variance *v*_*E*_ (resp. *v*_*I*_), such that each neuron receives the following recurrent input from each of the two populations *P* ∈ {*E, I*}:

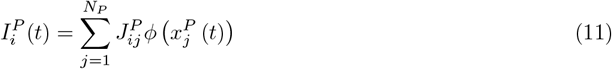

The mean and variance of the recurrent input affect the network dynamics differently. To account for this, we write the recurrent weights as a sum of their mean *m*_*P*_ and a random component 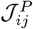 with mean 0 and variance 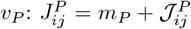 We further introduce the shorthand 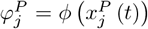, as well as the population-averaged firing rate: 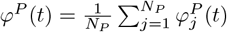. The recurrent input received by neuron *i* from population *P* is then:

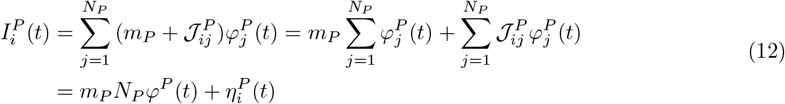

where

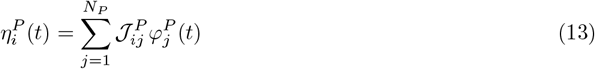

The total input received by each neuron *i* is then:

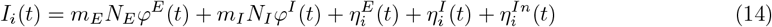

where 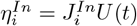.

### A.2 Statistics of the input to individual neurons

To determine the influence of the network connectivity on the statistics of the input received by each neuron, we model the input *I*_*i*_ received by each neuron as a Gaussian process of mean *µ* and variance Δ. We determine this mean and variance by averaging over the random connectivity, which we write as [·]. We note that, when averaging over the random connectivity, the statistics of the inputs received by all neurons are identical. As a consequence, the first and second-order moments of the population-averaged firing rates of the excitatory population match those of the inhibitory population, so we can write 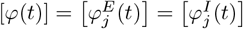 and 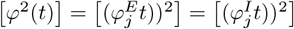.

#### A.2.1 Mean *µ* of the input to individual neurons

The mean input *µ*(*t*) = [*I*_*i*_(*t*)] is given by:

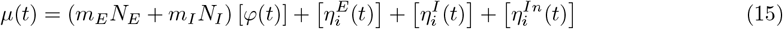

The input weights 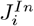 are independent of *U* (*t*) and have zero mean, therefore:

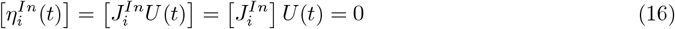

In the large network limit, the neuron firing rates are assumed to be independent of their outgoing weights, therefore 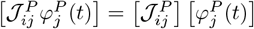. Since 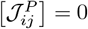, we have:

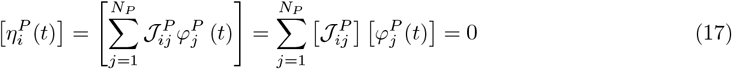

Therefore:

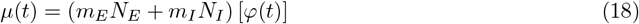

#### A.2.2 Variance Δ of the input to individual neurons

The variance Δ of the input to each neuron equals *I*_*i*_(*t*)^2^ −[*I*_*i*_(*t*)]^2^. Since the terms in the right-hand side of eq. (14) are all uncorrelated, the variance of *I*_*i*_ equals the sum of the variances of all terms. The variance of each of the first two terms equals 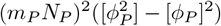 We assume that, in the large network limit, the population averages are identical to averages over the random connectivity: *φ*^*E*^(*t*) = *φ*^*I*^ (*t*) = [*φ*(*t*)]. With this assumption, these two variances will thus equal zero, and the recurrent input variance is equal to the variance of 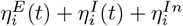. We decompose this into the following terms:

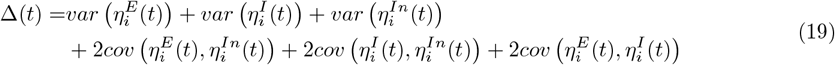

The variance of the external input is equal to:

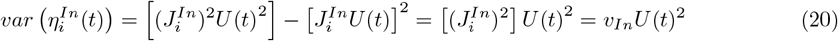

The external input weights 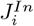 are assumed to be independent of the recurrent inputs, therefore:

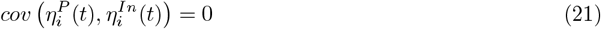

The variance of the recurrent inputs is equal to

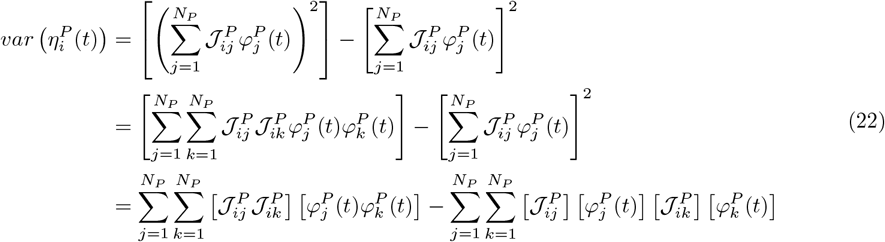

Since 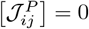, the last term is zero. Moreover, since the recurrent weights are independent:

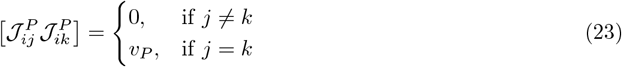

Therefore:

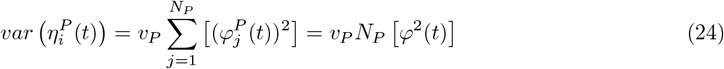

The covariance between the excitatory and inhibitory recurrent inputs is equal to:

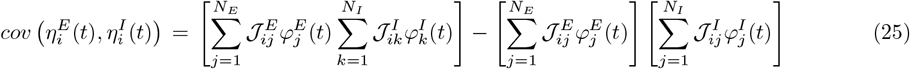

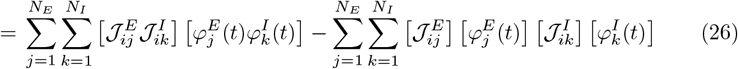

The recurrent weights from the two populations are independent, therefore 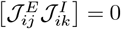 and:

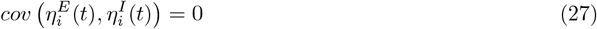

The input variance Δ is thus equal to:

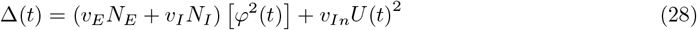

### A.3 Firing rate statistics

To obtain self-consistent equations for *µ*(*t*) and Δ(*t*), we need to express [*φ*(*t*)] and *φ*^2^(*t*) as functions of *µ*(*t*) and Δ(*t*). For this, we assume that the external input *U* (*t*) varies much more slowly than the time constant *τ*, and that the system is in a quasi-stationary state. We will show below (see section A.6) that this stationary state is stable. For stationary solutions, the activation *x*_*i*_(*t*) has the same distribution as the input *I*_*i*_(*t*). We explicitly model the input *x* as a Gaussian process whose mean *µ*(*t*) and variance Δ(*t*) satisfy eqs. (18) and (28): 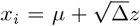, where *z* is a Gaussian process with zero mean and unit variance, whose probability distribution is thus given by 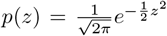. By modelling the input as a Gaussian, we assume that the input has zero skewness. Simulation results show that the skewness is indeed small for all investigated imbalances (fig. 8).

**Fig. 8.**
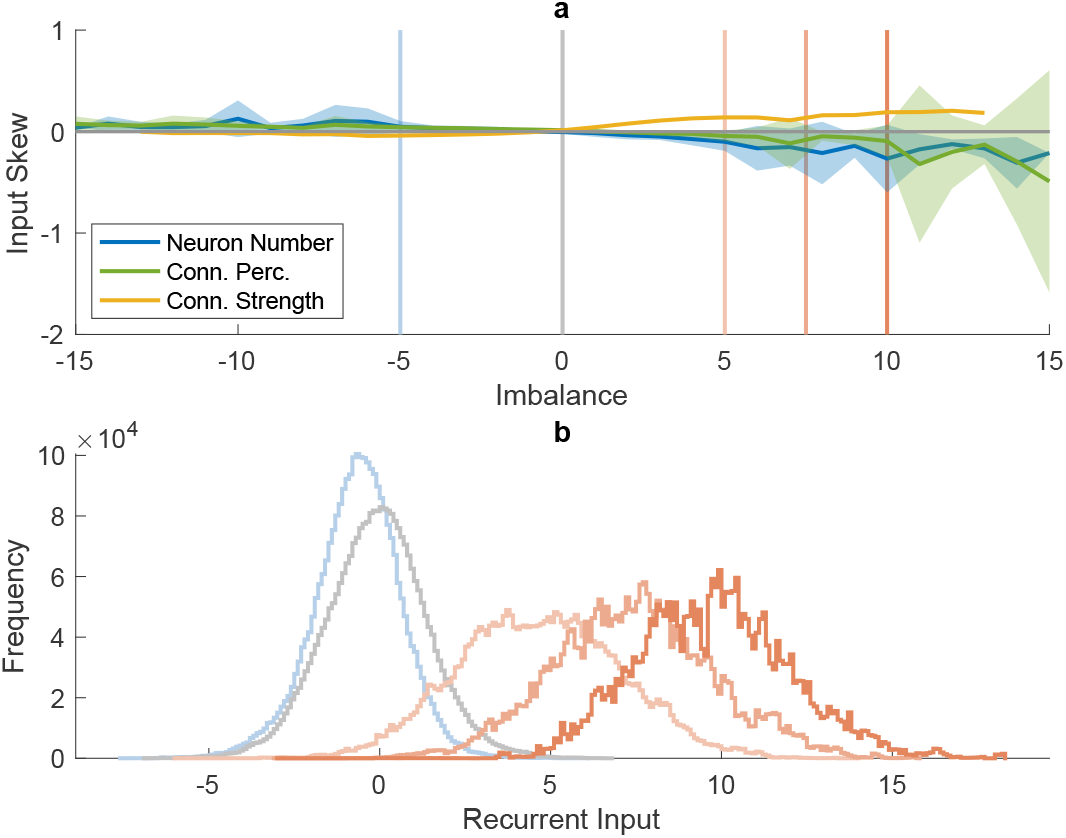
Skew of the time-averaged recurrent input distributions. (a) shows the input skew as a function of the imbalance *A* when varying the neuron number (blue), connection percentage (green) and connection strength (yellow), as well as the corresponding MFT predictions (black) for *v*_*E*_ *N*_*E*_ + *v*_*I*_ *N*_*I*_ = 2 *·* 1.5^2^. For the neuron number and connection percentage, the average and standard deviations over all levels of *N*_*tot*_ and *p*_*tot*_ are shown. For the connection strength, only *v*_*E*_ *N*_*E*_ + *v*_*I*_ *N*_*I*_ = 2· 1.5^2^ is shown to match MFT predictions. The coloured vertical lines indicate the imbalance levels of the four example networks shown in panel (b). (b) shows the histograms of the recurrent input in these four networks

We can then obtain the firing rate statistics by averaging over *z*:

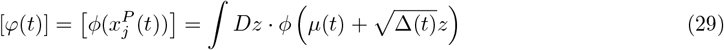

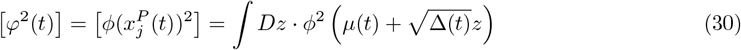

where 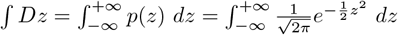

### A.4 Solving the equations

Substituting eqs. (18) and (28) into eqs. (29) and (30), we obtain the following equations for the *µ*- and Δ-nullclines, N_*µ*_ and N_Δ_:

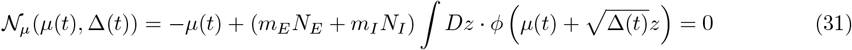

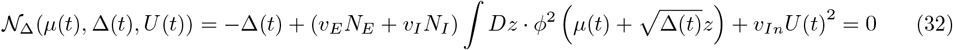

We solved the system of equations numerically using a built-in solver for sets of non-linear equations, fsolve, in Matlab R2023a (MathWorks, Inc., Natick, Massachusetts, United States). Graphically, this corresponds to finding the intersection between the *µ*- and Δ-nullclines.

#### A.4.1 Nullclines

The *µ*-nullcline depends on the imbalance *m*_*E*_*N*_*E*_ + *m*_*I*_ *N*_*I*_. It is illustrated in fig. 9 for different values of imbalance ranging from -10 to 10. For large, positive imbalance (red curves in fig. 9) the *µ*-nullcline approaches a vertical line at *µ* = *m*_*E*_*N*_*E*_ + *m*_*I*_ *N*_*I*_. In contrast, for large, negative imbalance (blue curves in fig. 9), *µ* decreases only moderately and does not scale with *m*_*E*_*N*_*E*_ + *m*_*I*_ *N*_*I*_.

**Fig. 9.**
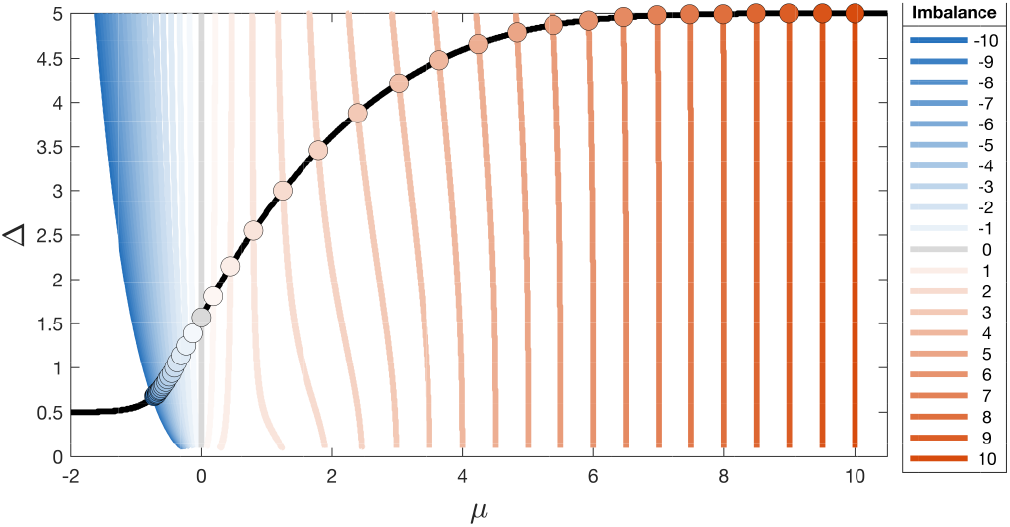
Graphical representation of the *µ* and Δ nullclines. The *µ* nullcline (coloured) depends on the anatomical imbalance *A*, shown here for values ranging from -10 to 10. The Δ nullcline (black) depends on *v*_*E*_*N*_*E*_ + *v*_*I*_ *N*_*I*_, shown here in black for a single value 2 · 1.5^2^, as used in the simulations. The solutions for each imbalance level, indicated by the coloured dots, are found at the intersection of the two nullclines

The Δ-nullcline depends on the recurrent weight variance *v*_*E*_*N*_*E*_ + *v*_*I*_ *N*_*I*_ and the external input variance *v*_*In*_*U* ^2^(*t*). It is illustrated in fig. 9 for a recurrent weight variance of *v*_*E*_*N*_*E*_ + *v*_*I*_ *N*_*I*_ = 2 ·1.5^2^ and *v*_*In*_*U* ^2^(*t*) = 1*/*2 (as used in simulations throughout the paper, unless otherwise specified). The variance Δ decreases with decreasing *µ* until a minimum value Δ_*min*_ = *v*_*In*_*U* (*t*)^2^. For increasing *µ*, it increases up to a plateau value of Δ_*max*_ = *v*_*In*_*U* ^2^(*t*) + *v*_*E*_*N*_*E*_ + *v*_*I*_ *N*_*I*_.

The solutions for *µ* (fig. 10a) and Δ (fig. 10b) are given by the intersection of the two nullclines. The obtained values are subsequently used to calculate the firing rate mean (fig. 10c) and variance (fig. 10d) using eqs. (29) and (30).

**Fig. 10.**
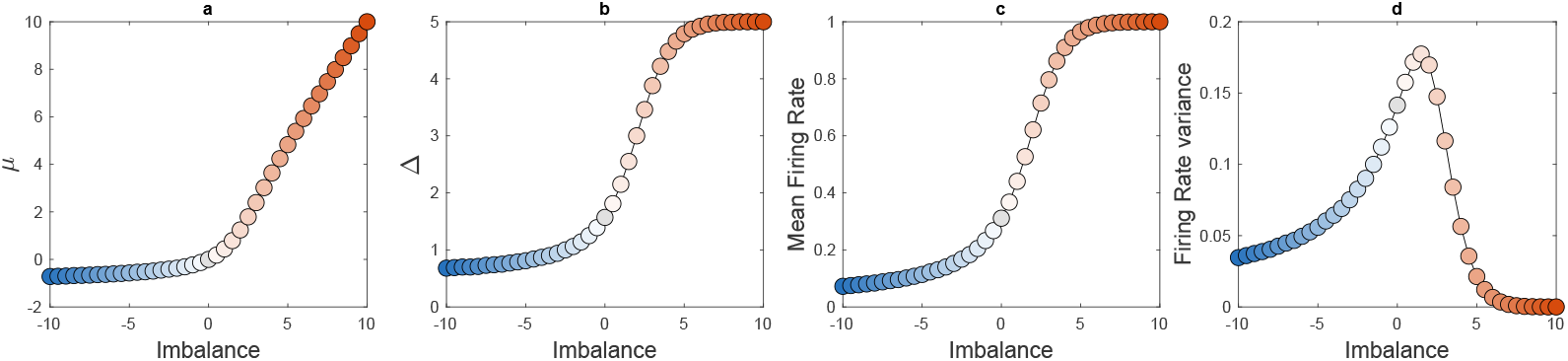
Mean-field theory solutions found via the nullcline method for *v*_*E*_*N*_*E*_ + *v*_*I*_ *N*_*I*_ = 2 · 1.5^2^ and a rectified hyperbolic tangent as activation function, as used in the simulations. (a) shows the mean recurrent input, (b) the recurrent input variance, (c) the mean firing rate, and (d) the firing rate variance as a function of the imbalance *A*

The mean-field theory equations we derived are valid for any transfer function, so our results can easily be applied to different networks. Using a sigmoid transfer function instead of the rectified hyperbolic tangent, would have led to similar results (see fig. 11) The largest difference can be seen in the firing rate variance, which is lower when using a sigmoid (cf. panel D in figs. 10 and 11).

**Fig. 11.**
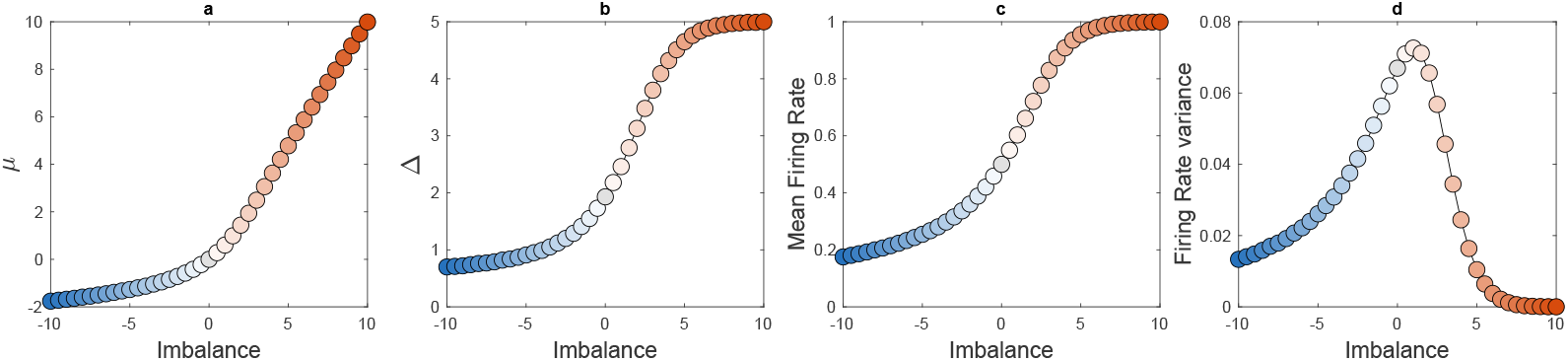
Mean-field theory solutions found via the nullcline method for networks using a sigmoid activation function with *v*_*E*_*N*_*E*_ + *v*_*I*_ *N*_*I*_ = 2 · 1.5^2^. (a) shows the mean recurrent input, (b) the recurrent input variance, (c) the mean firing rate, and (d) the firing rate variance as a function of the imbalance *A*

### A.5 Mean and variance of the weight distributions

In our simulations, the outgoing connection weights from population *P* have a probability *p*_*P*_ of being non-zero. The non-zero weights are drawn from a Gaussian distribution with mean 0 and variance 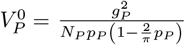. This scaling is chosen such that the variance of the weight distribution scales as 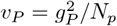 (see below). For excitatory weights, the absolute value of this weight is taken, whereas for inhibitory weights, the negative of the absolute value is taken. The mean *M*_*P*_ and variance *V*_*P*_ of the non-zero weights, therefore, correspond to those of a half-Gaussian distribution:

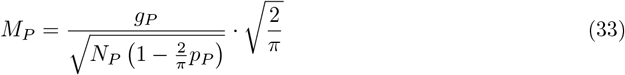

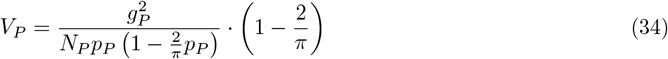

The mean recurrent weight of the full connection matrix *m*_*P*_ is therefore equal to

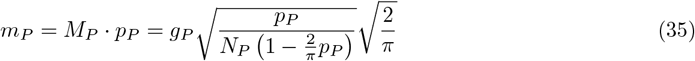

To obtain the variance of the full matrix, we can do the same for the mean of the squared connection weight, which equals 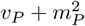, to get:

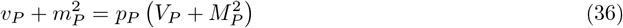

Therefore:

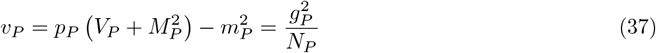

We then obtain the mean and variance of the sum of the recurrent weights for our networks:

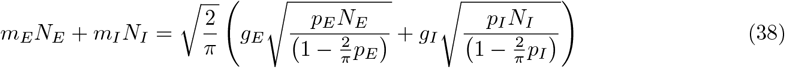

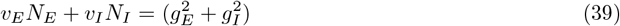

Here, *m*_*E*_*N*_*E*_ + *m*_*I*_ *N*_*I*_ corresponds to the anatomical imbalance *A*, and *v*_*E*_*N*_*E*_ + *v*_*I*_ *N*_*I*_ is equal to the square of the total connection strength *g*_*tot*_, as defined in the main text.

The input weight matrix was taken from a Gaussian distribution with mean zero and variance 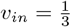. The external input *U* (*t*) is a sinusoid ranging from 0 to 2. As we analysed a quasi-stationary state, the MFT was performed using a single value for the external input, corresponding to the temporal mean of *U* : ∫ *U* (*t*)^2^ = 1.5. Therefore, the term *v*_*in*_*U* (*t*)^2^ equalled 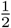 in all included MFT analyses.

### A.6 Stability Analysis

#### A.6.1 Local perturbation analysis

To determine whether the stationary solutions derived above are stable, we linearized the system dynamics around the steady state. To simplify notations, we introduce the *N*_*tot*_ by *N*_*tot*_ (with *N*_*tot*_ = *N*_*E*_ + *N*_*I*_) matrix ***J*** whose first *N*_*E*_ columns correspond to the excitatory weights 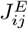 and whose last *N*_*I*_ columns correspond to the inhibitory weights 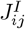. The system dynamics are then:

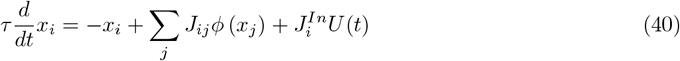

and the steady-state 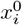 satisfies:

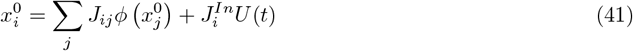

To linearize the dynamics, we consider small perturbations *δx*_*i*_ around the steady state 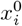 and write 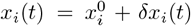, where 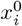 is constant. From this, it follows that 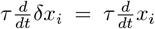. We can then approximate the derivative of *δx*_*i*_ using a first-order Taylor series expansion to get:

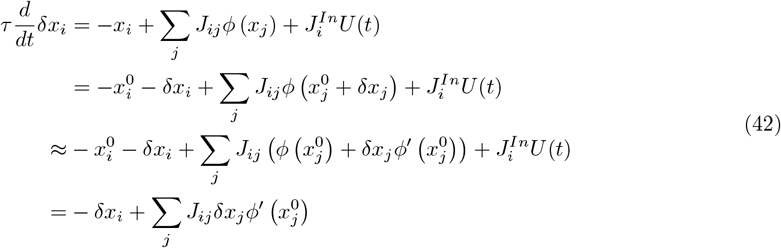

We can write this in matrix notation, using the perturbation vector 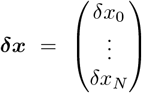, the *N* _*tot*_-dimensional identity matrix ***I*** and 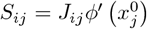, resulting in:

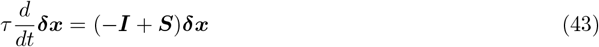

This system is stable if and only if the eigenvalues of (− ***I*** + ***S***) all have a negative real part or, equivalently, if the eigenvalues of ***S*** all have real parts smaller than 1. The eigenvalue spectrum of ***S*** consists of *N*_*tot*_ − 1 eigenvalues distributed within a disk in the complex plane, and a single outlier eigenvalue (Rajan and Abbott, 2006; Mastrogiuseppe and Ostojic, 2018), as shown in fig. 12a. The disk corresponds to the network’s response to local perturbations to individual neurons, while the outlier eigenvalue corresponds to the network’s response to a global perturbation of the population-averaged mean *µ* or variance Δ of the input.

The radius *r* of the disk can be calculated by summing the variances of the columns of ***S***:

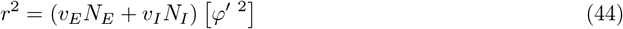

When this radius is larger than unity, the stationary state loses stability and the network has chaotic, fluctuating activity (Rajan and Abbott, 2006; Aljadeff et al., 201<u>5</u>; Harish and Hansel, 2015; Mastrogiuseppe and Ostojic, 2018). We find that this occurs when 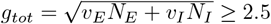 (see fig. 12b), and exclude these networks from the results.

**Fig. 12.**
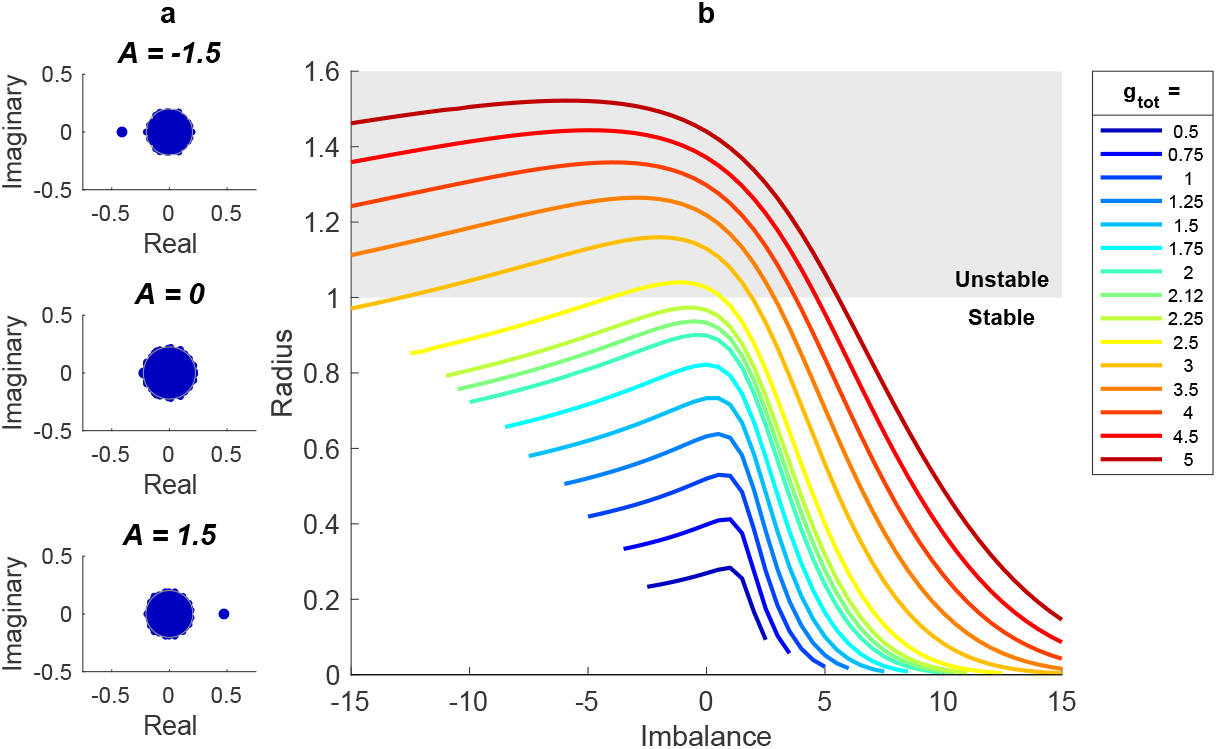
Results of the local stability analysis. (a) shows three example eigenvalue spectra from simulated networks with total connection strength *g*_*tot*_ = 0.5 and imbalance *A* = *−*1.5, 0 and 1.5. (b) shows the MFT-derived radius of the eigenvalue spectrum of the stability matrix as a function of the imbalance *A* for various levels of the total connection strength 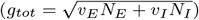. The grey-shaded area indicates radii for which the network is unstable

#### A.6.2 Global perturbation analysis

It is not known how to calculate the outlier eigenvalue directly, but following the approach of Kadmon and Sompolinsky (2015) and Mastrogiuseppe and Ostojic (2018), we derive it by assessing the response of the population-averaged mean *µ* and variance Δ to a global perturbation. First, we derive a dynamical system governing the temporal evolution of *µ* and Δ. Then, we linearize this system around the steady state *µ*_0_, Δ_0_ obtained by solving eqs. (31) and (32).

##### *Time-evolution of µ and* Δ

We extend the previous mean-field theory analysis to consider *µ*(*t*) = [*I*_*i*_(*t*)], i.e. the average (across different instantiations of the random connectivity) of the input received by neuron *i* after a global perturbation occurring at time *t* = 0. We decompose the neural activation response *x*_*i*_(*t*) as a sum of this population-averaged response *µ*(*t*) and individual neuron variation *ξ*_*i*_(*t*):

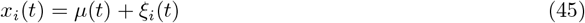

The dynamics of *x*_*i*_ can therefore be written as:

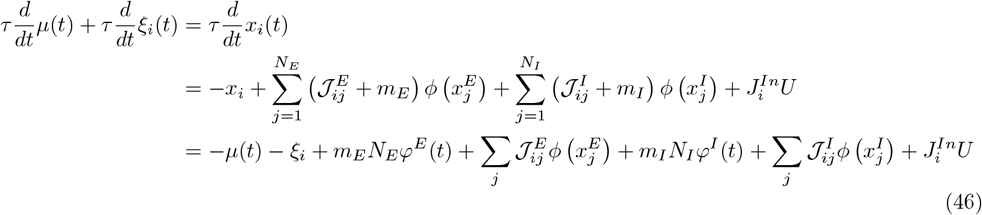

We then average over the random connectivity, substituting 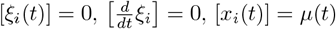 and 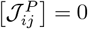 in the above equation, to obtain the following differential equation for *µ*:

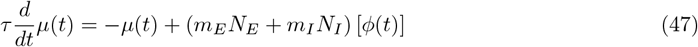

To obtain a differential equation of Δ(*t*) = *ξ*_*i*_(*t*)^2^, we subtract eq. (47) from eq. (46). As we did previously, we consider that, in the large network limit, the population averages are identical to averages over the random connectivity: *φ*^*E*^(*t*) = *φ*^*I*^ (*t*) = [*φ*(*t*)]. We then obtain:

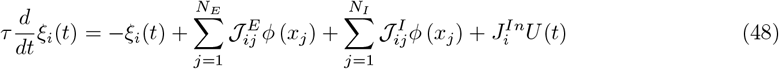

We note that 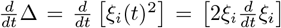 and use this to express 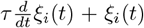 in the formula above in terms of Δ(*t*) and 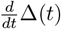:

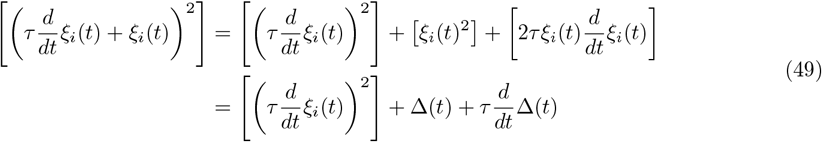

As temporal derivatives vanish when evaluated at fixed points, we can neglect the 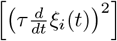 term. Substituting equation eq. (48) into eq. (49), we obtain:

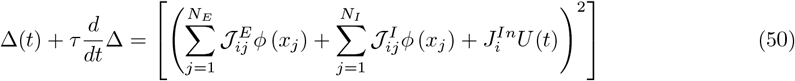

The right-hand term can be evaluated as in section A.2, resulting in:

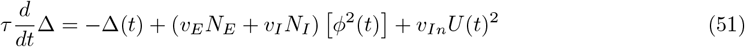

##### Linearization

We obtain the dynamics of the system’s response to global perturbations by substituting the equations for the *µ*- and Δ-nullclines (eqs. (31) and (32)) into eqs. (47) and (51):

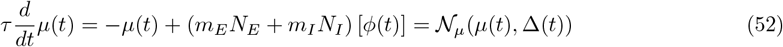

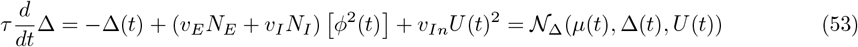

To determine whether the system is stable, we linearize the system dynamics around the steady-state *µ*_0_, Δ_0_: *µ*(*t*) = *µ*_0_ + *δµ*(*t*),Δ(*t*) = Δ_0_ + *δ*Δ(*t*).

The response of the mean input follows:

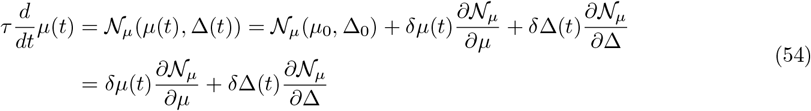

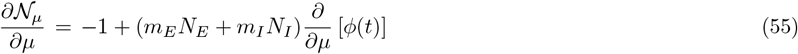

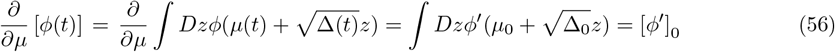

where [.]_0_ denotes the average evaluated at the steady-state.

For the variance, we get:

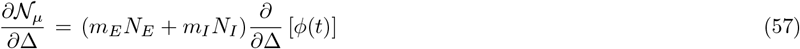

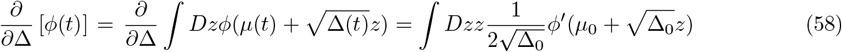

We use ∫ *Dz zf* (*z*) = ∫ *Dz f*^*′*^(*z*) (derived below in section A.6.3). Substituting this into the equation above, we get:

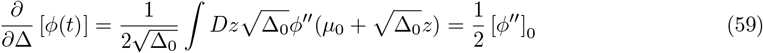

Substituting into eq. (54), we obtain:

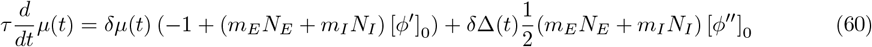

We similarly obtain the linearized dynamics for Δ(*t*):

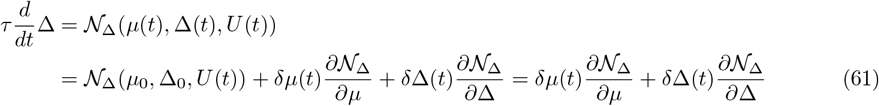

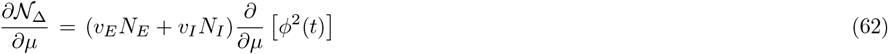

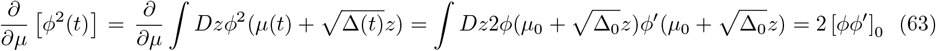

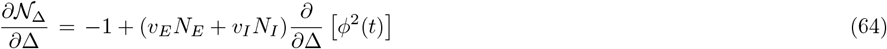

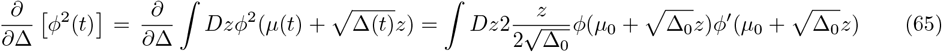

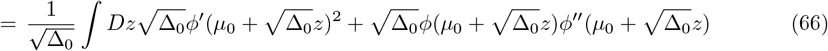

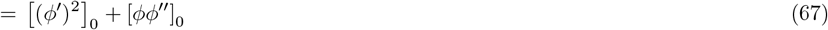

Substituting this into eq. (61), we obtain:

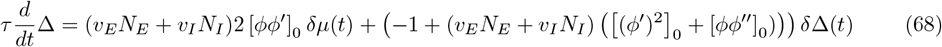

Combining eq. (60) and eq. (68), we obtain:

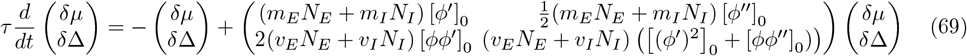

The system is stable to global perturbations if and only if the eigenvalues of ***M*** have real part smaller than 1, where:

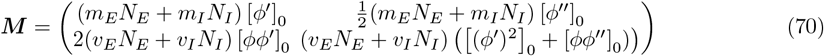

To simplify notations, we introduce:

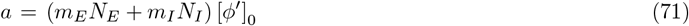

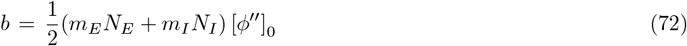

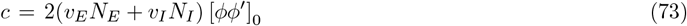

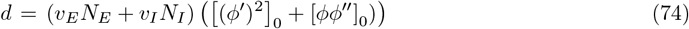

The eigenvalues of ***M*** are the roots of the following polynomial:

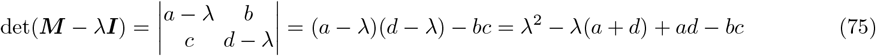

The discriminant of this polynomial is:

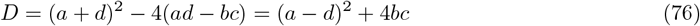

The roots *λ*_+_, *λ*_*−*_ are thus given by:

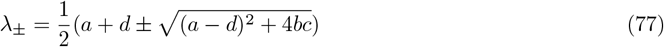

These roots are illustrated in fig. 13. Note that, when *m*_*E*_*N*_*E*_ + *m*_*I*_ *N*_*I*_ is close to zero, both roots are smaller than the radius of the disk of eigenvalues corresponding to the system’s response to local perturbations (equation eq. (44)). In this case, the matrix ***S*** does not have an outlier eigenvalue (see fig. 12a, middle). For positive *m*_*E*_*N*_*E*_ + *m*_*I*_ *N*_*I*_ and small *v*_*E*_*N*_*E*_ + *v*_*I*_ *N*_*I*_, we have *λ*_+_ *> r* (and |*λ*_*−*_| *< r*), and ***S*** has one outlier eigenvalue equal to *λ*_+_ (see fig. 12a, bottom, and fig. 13). For negative and large enough *m*_*E*_*N*_*E*_ + *m*_*I*_ *N*_*I*_, we have *λ*_*−*_ *<* −*r* (and |*λ*_+_| *< r*), and ***S*** has one outlier eigenvalue equal to *λ*_*−*_ (see fig. 12a, top and fig. 13). The real parts of both *λ*_+_ and *λ*_*−*_ are always smaller than 1 (see fig. 14), so the system is always stable to global perturbations and its stability will only depend on the radius (local stability).

**Fig. 13.**
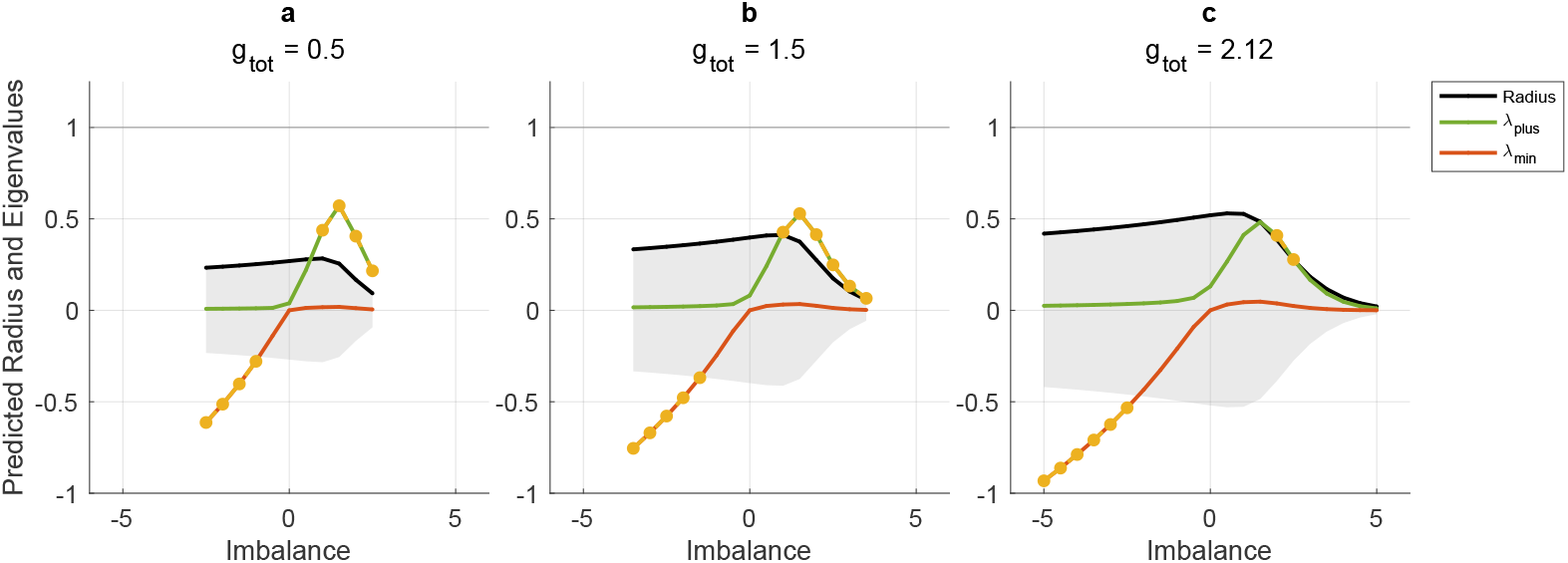
Example results of the global stability analysis. The predictions of the radius and outliers of the global stability matrix are shown for three levels of total connection strength (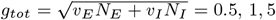, and 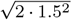). Each panel shows the positive (*λ*_+_, green line) and negative (*λ*_*−*_, red line) eigenvalues of the global stability matrix ***M***, as well as the radius of its eigenvalue spectrum (black line). The grey area indicates where the eigenvalues are smaller than the radius of the eigenvalue spectrum. Yellow dots indicate outliers, i.e., maximum eigenvalues that are larger than the radius

**Fig. 14.**
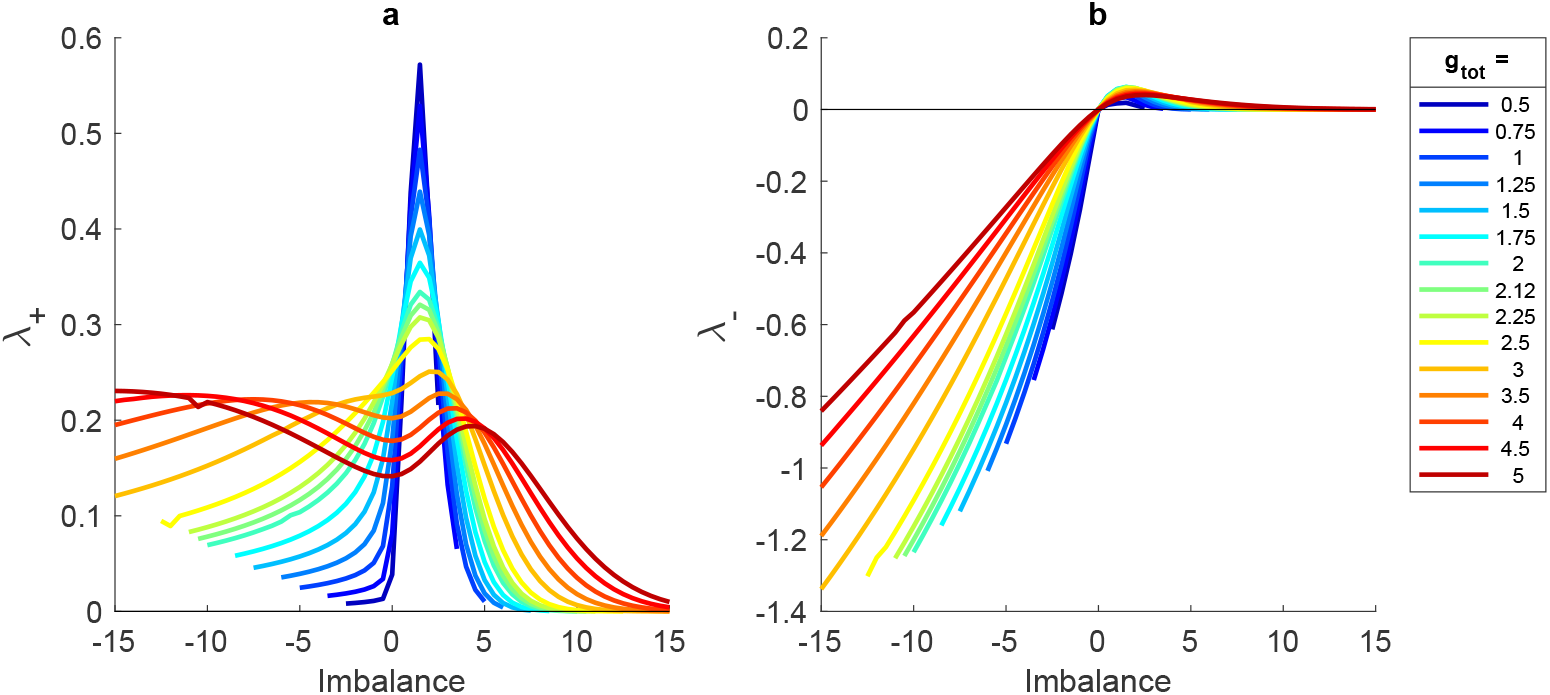
Results of the global stability analysis. Mean-field theory derived values of the (a) positive (*λ*_+_) and (b) negative (*λ*_*−*_) eigenvalues of the global stability matrix ***M*** are shown as a function of the imbalance *A*, for various level of the total connection strength 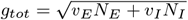

#### A.6.3 Integration by parts

We consider any function *f* (*z*) such that 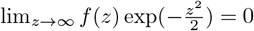:

We introduce :

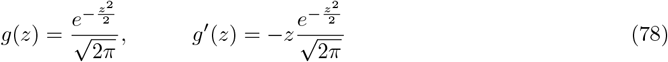

We have:

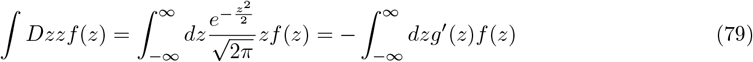

Using integration by parts, we get:

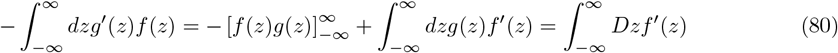

Thus:

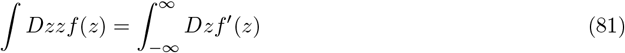

## Appendix B Network training algorithm

To train the neural networks, we use a recursive least squares algorithm that updates the output weights at each time step, as described by Haykin (2014) and employed in the FORCE algorithm of Sussillo and Abbott (2009).

The output signals of the network were calculated by multiplying the [*k* ×*N*]-dimensional output weights matrix ***W*** (*t*) with the sigmoid-transformed neuron activations ***φ***(*t*) = *ϕ*(***x***(*t*)).

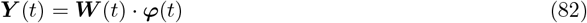

The output error ***ϵ***, a *k*-dimensional row vector, was calculated as the difference between the calculated output ***Y*** (*t*) and the target output ***Y***_***target***_(*t*).

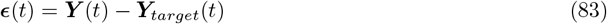

The output weights ***W*** (*t*) were updated based on the current activity, the output error ***ϵ***, and the [*N*_*tot*_ × *N*_*tot*_] learning rate matrix ***P*** (*t*). Here, ***P*** (*t*) denotes an estimate of the inverse of the correlation matrix of ***φ***(*t*) with a regularization term (Sussillo and Abbott, 2009).

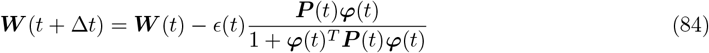

***P*** (*t*) was updated according to:

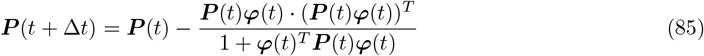

## Appendix C

Calculation of spinal cord efferent and afferent signals using computational musculo-skeletal model Myonardo

*Myonardo* is a Matlab-based, 3D computational musculo-skeletal model developed in the Simscape Multibody 3D simulation environment, consisting of 23 bone segments, 23 joints and 180 muscle-tendon-units (Predimo GmbH, Münster, Germany). The relative mass and size of the individual segments, as well as the muscular attachments were determined based on the work of Shippen and May (2012). The model was scaled to the subject’s body mass and height (Winter, 2009; Hatze, 1980).

As a first step, *Myonardo* calculated the muscle-tendon lengths, muscle-tendon velocities and the muscular lever arms relative to the instantaneous joint centre, based on the instantaneous joint angles and joint angular velocities acquired from the 3D kinematics. The muscle line of action between the origin and the insertion of each muscle-tendon unit is separated into muscle parts between via-points, such that the muscle length was calculated as the sum of the length of each muscle part, i.e. the Euclidean distance between the attachments of the muscle part.

Based on these muscle lengths *l*_*m*_ and velocities *v*_*m*_, the maximum possible muscle force *f*_*m,max*_, could then be calculated using the force-velocity relationship (*f*_*v*_), as well as the active (*f*_*la*_) and passive (*f*_*lp*_) force-length relationships as follows:

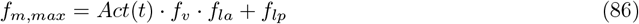

where *Act*(*t*), the activation of the muscle in the interval [0,1], was set to 1. The Hill-type force-velocity relationship (*f*_*v*_) was calculated as:

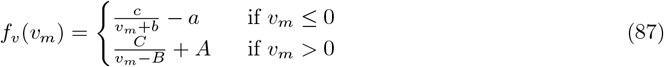

where properties *a, b, c* and *A, B, C* were estimated from the isometric force *f*_*iso*_, resting length *l*_*opt*_, eccentric gain *f*_*ecc*_ and the ratio of fast-twitch to slow-twitch fibres *FT* %, using parameter values from Thaller and Wagner (2004) and Herzog (1999). The active (*f*_*la*_) and passive (*f*_*lp*_) force-length relations were calculated as:

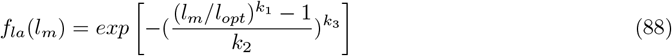

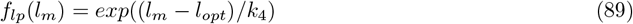

with *k*_1_ = 0.96, *k*_2_ = 0.35, *k*_3_ = 2, *f*_*ecc*_ = 1.5, and *FT* % = 0.5. The remaining muscle properties *f*_*iso*_ and *l*_*opt*_ were taken from Rajagopal et al. (2016).

Based on the maximum force *f*_*m,max*_ for each muscle and the lever arms *r*, the maximum torque that each muscle can generate around a joint at a certain instance of time was calculated as *T*_*max*_ = *r*× *f*_*m,max*_. Then, a set of muscular activations *Act*(*t*), within the boundary 0 ≤ *Act* ≤ 1, can be determined for each joint, such that

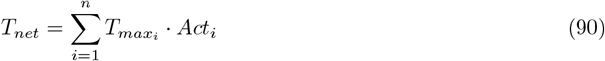

with *i* indicating the *n* muscles that are acting at the given joint. The required net torque *T*_*net*_ is known from inverse dynamics and the maximum torque *T*_*max*_ was calculated as described above. As there are more muscles than degrees of freedom of a joint contributing to the joint-torques, the muscular activations of these redundant muscles have to be distributed via optimization. In the present study, the muscular activation was calculated by minimizing the sum of squared muscular activations for each instant of time-based on the linear least-squares solver (lsqlin):

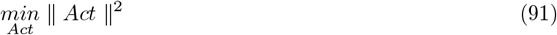

## Appendix D Supplementary Figures

**Fig. 15.**
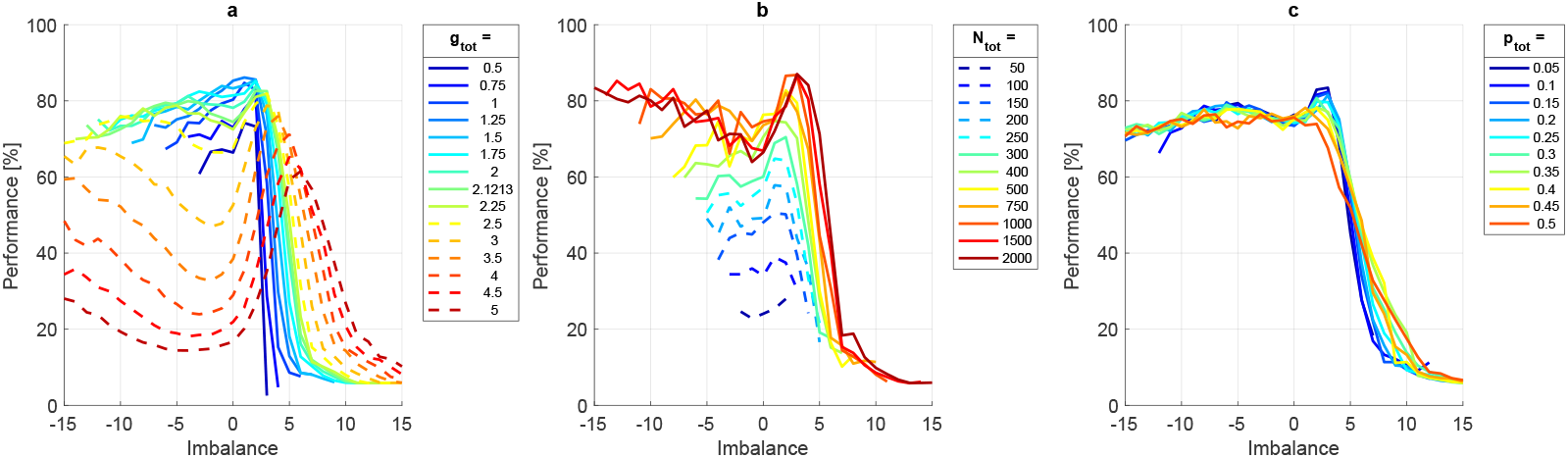
Performance as a function of the imbalance. *A*, for all investigated levels of (a) *g*_*tot*_, (b) *N*_*tot*_, and (c) *p*_*tot*_. Dashed lines indicated networks that have been excluded from the main results (i.e., those with *N*_*tot*_ *≤* 250 and *g*_*tot*_ *≥* 2.5)

**Fig. 16.**
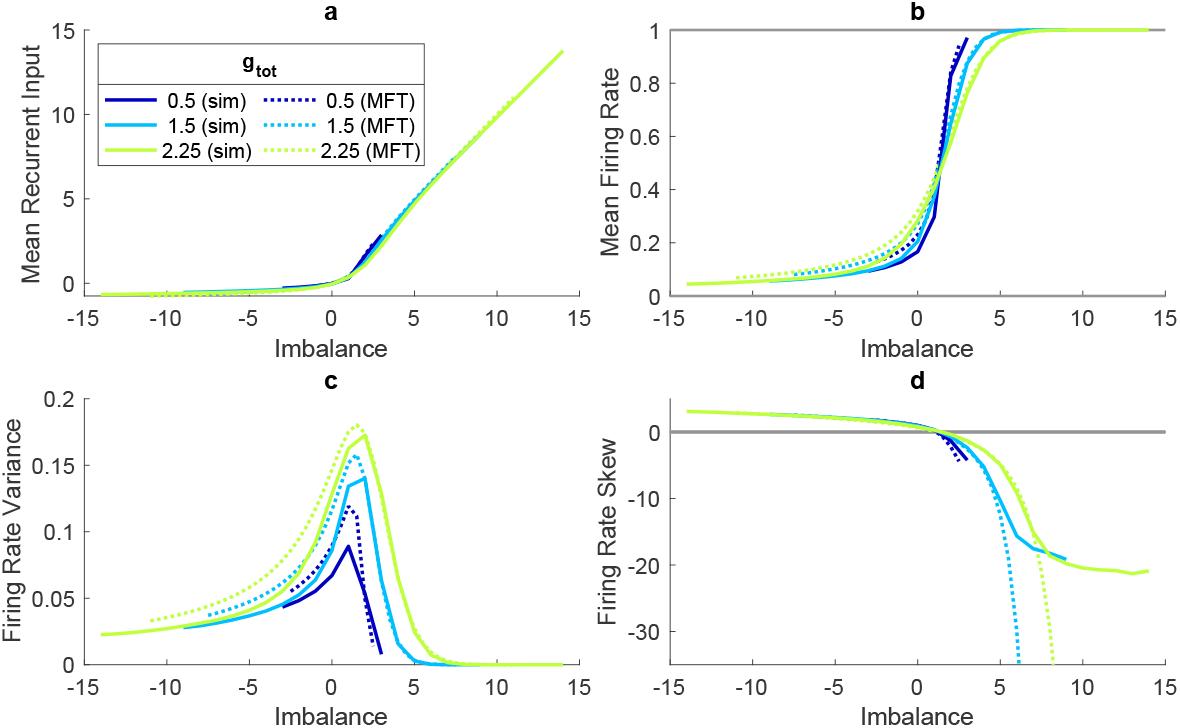
Mean-field theory predictions (dotted)versus simulation results (solid) for three levels of total connection strength (*g*_*tot*_ = 0.5, 1.5, and 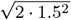). Note that, for the simulations, results without output feedback are shown, as output feedback has not been taken into account in the MFT predictions

**Fig. 17.**
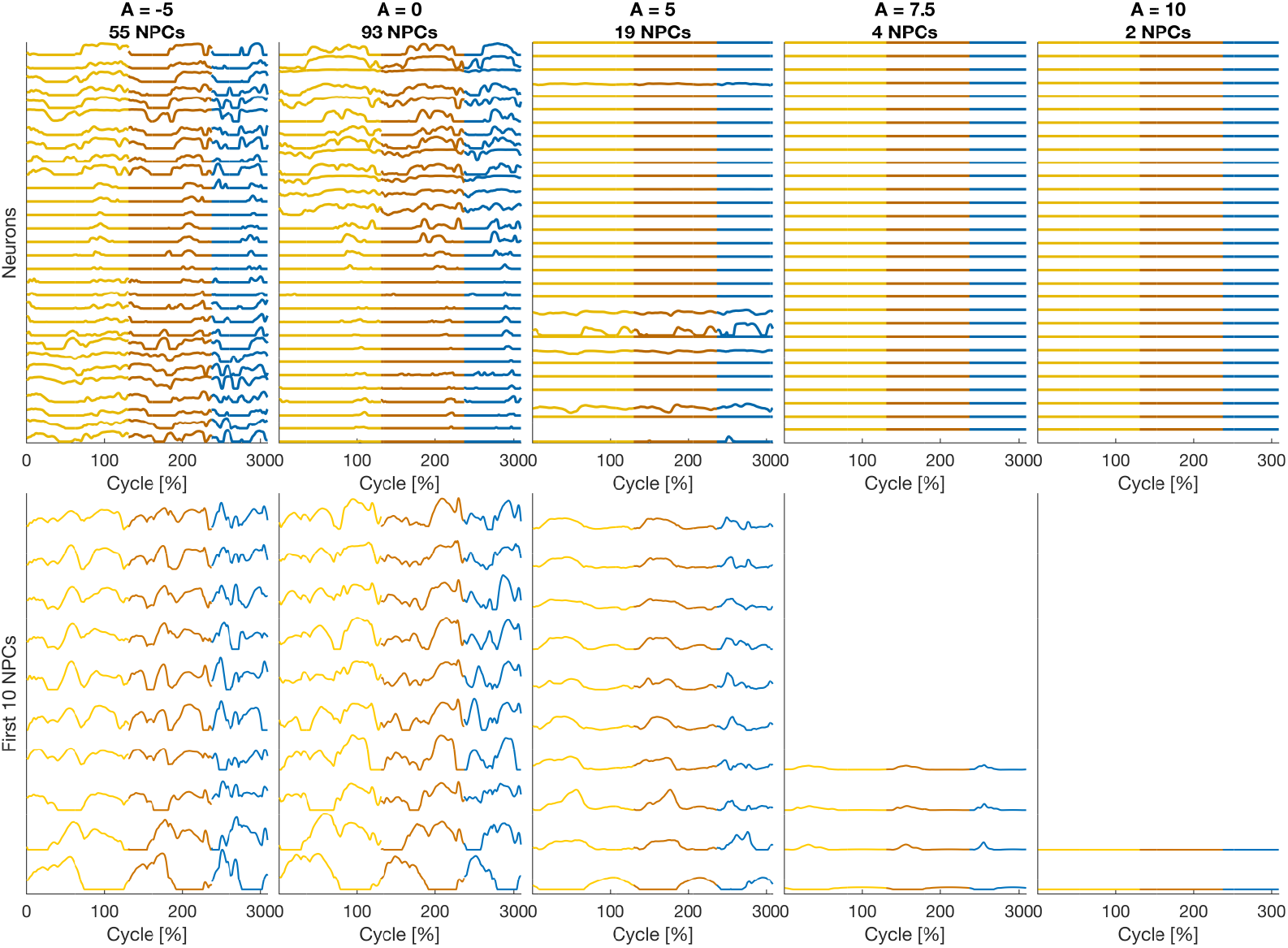
Example neural firing rates (*φ*) and resulting network principal components (NPCs) for five example networks with imbalances of *A* = *−* 5, 0, 5, 7.5 and 10 from left to right. The top row displays the firing rates for a selection of 30 out of 750 neurons. The bottom row shows up to ten NPCs resulting from the full set of neural activity patterns. The NPCs are ordered by their contribution to the overall variance, with those explaining the largest variance at the bottom. One example cycle for each of the three locomotor activities is illustrated, with, from left to right, walking slowly (yellow), walking fast (orange) and running (blue)

**Fig. 18.**
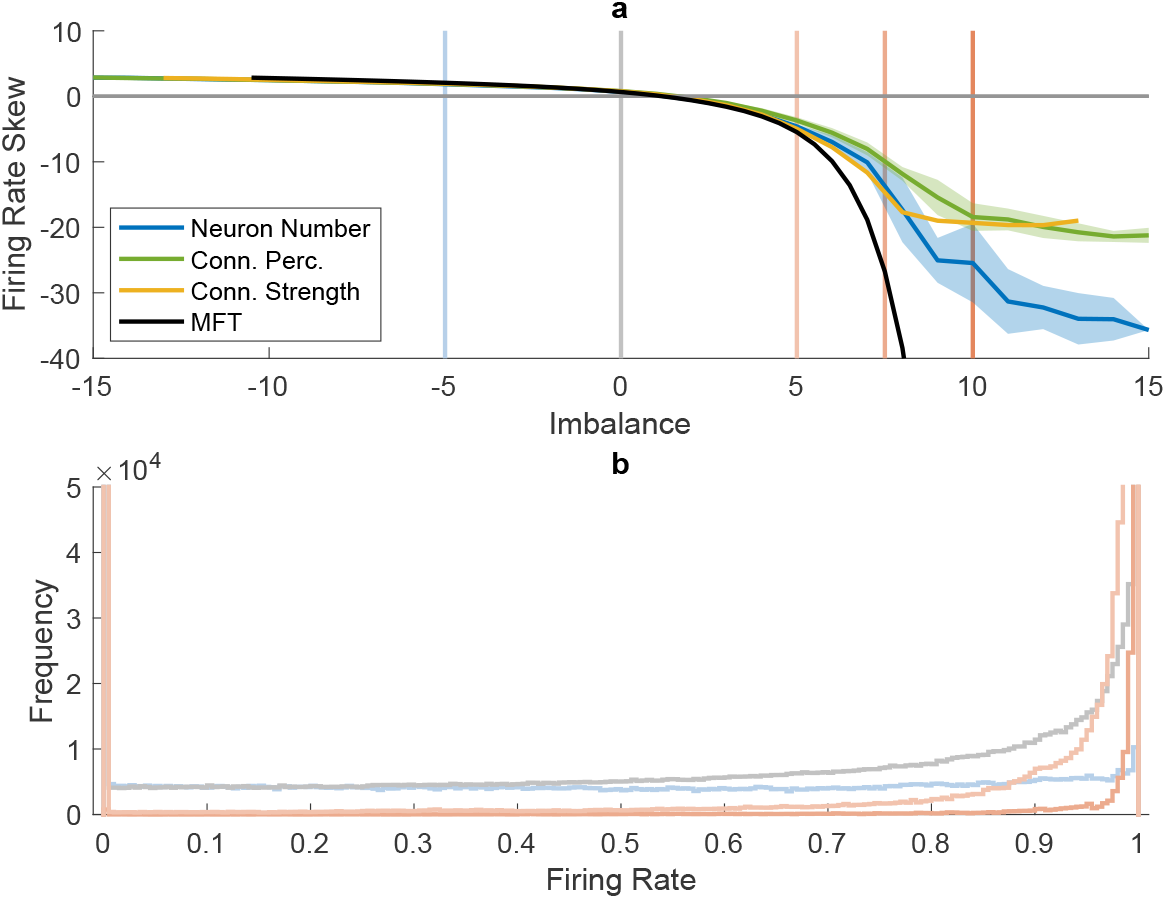
Skew of the time-averaged firing rate distributions. (a) shows the firing rate skew as a function of the imbalance (*A*) when varying the neuron number (blue), connection percentage (green) and connection strength (yellow), as well as the corresponding MFT predictions for 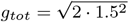. For the neuron number and connection percentage, the average and standard deviations over all levels of *N*_*tot*_ and *p*_*tot*_ are shown. For the connection strength, only 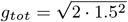 is shown, to match MFT predictions. The coloured vertical lines indicate the imbalance levels of the four example networks shown in panel (b). (b) shows the firing rate histograms of these four networks, zoomed in to the (0 *−* 5*e*^4^) range

**Fig. 19.**
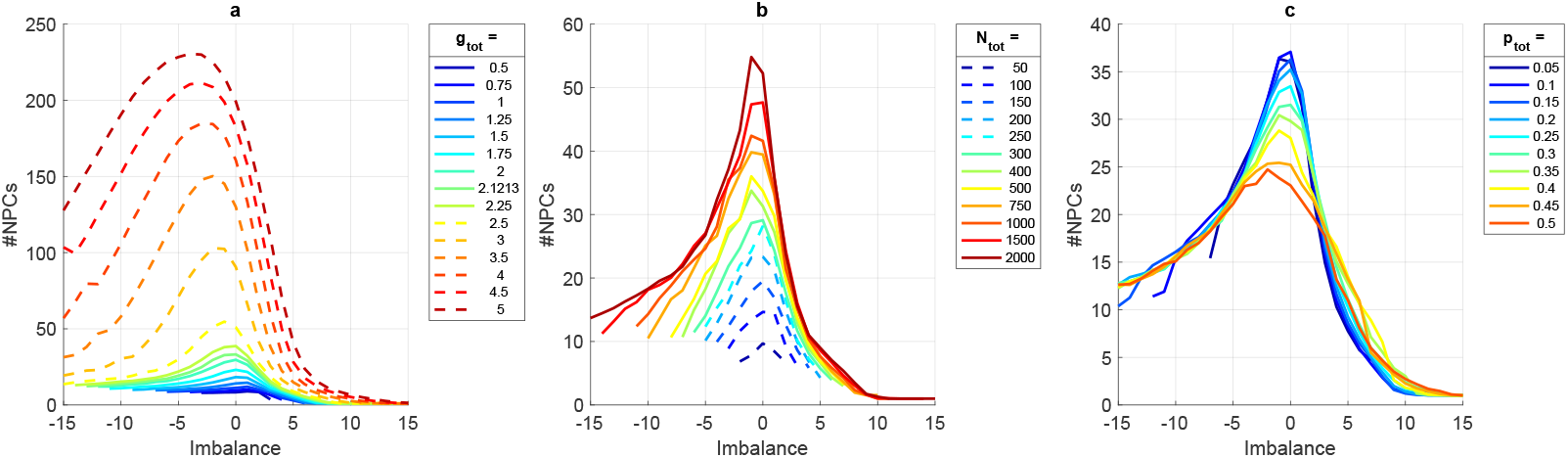
Number of principle components (NPCs) as a function of the imbalance. *A*, for all investigated levels of (a) *g*_*tot*_, (b) *N*_*tot*_, and (c) *p*_*tot*_. Dashed lines indicated networks that have been excluded from the main results (i.e., those with *N*_*tot*_ *≤* 250 and *g*_*tot*_ *≥* 2.5)

**Fig. 20.**
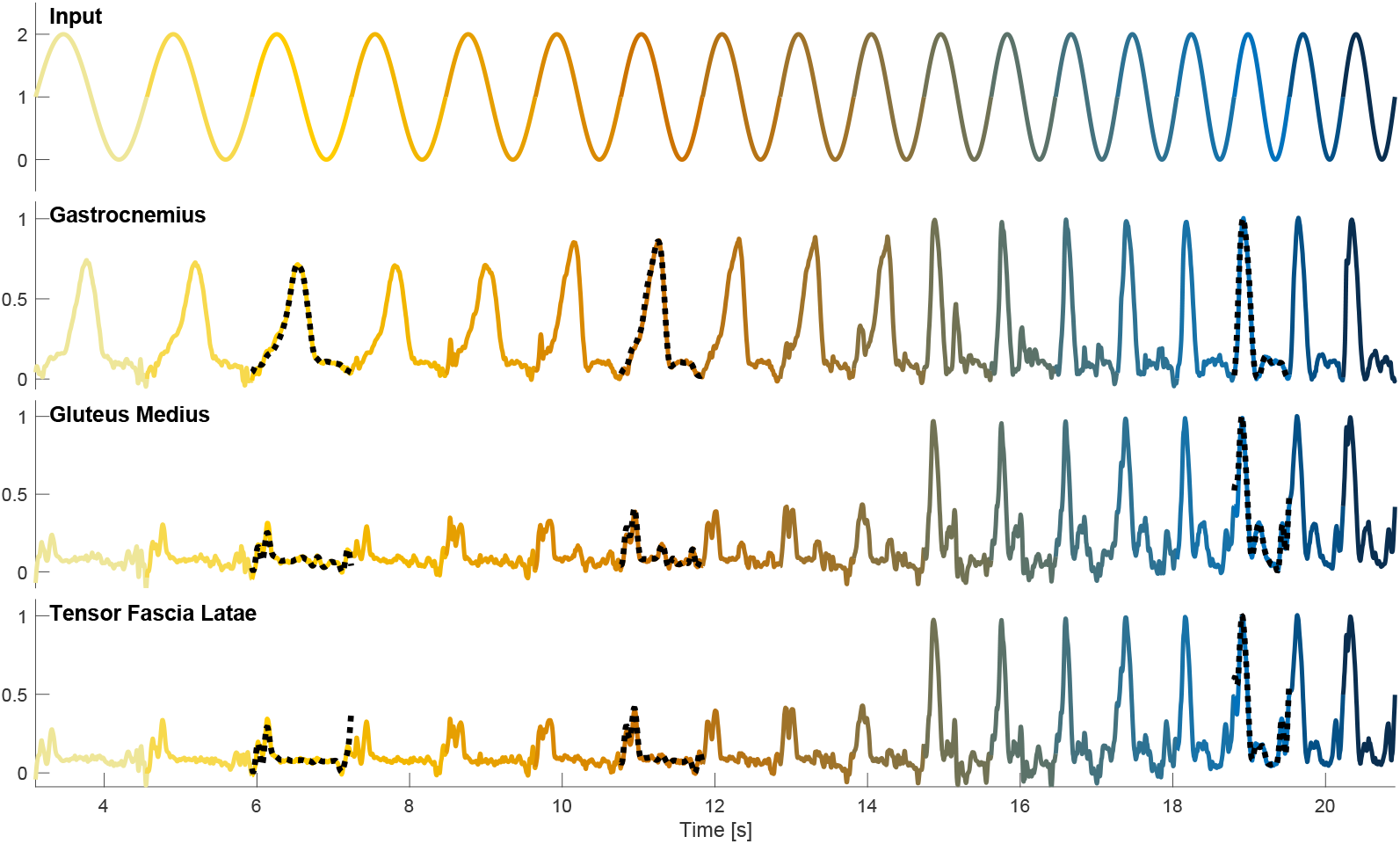
Example outputs for a morphing task. An example network with imbalance *A* = 0 was trained normally and tested on a morphed signal, where each cycle had a higher frequency than the one before. The top row shows the input signal, the other rows show network outputs for three example muscles: m. gastrocnemius, m.gluteus medius and m. tensor fascia latae. The colour of the signals reflects the frequency of each cycle (with yellow for slow walking, orange for fast walking, and blue for running). Output targets for the trained cycles are overlaid as dotted black lines. Output targets for the morphed cycles were not available. The example network had *N*_*E*_ = *N*_*I*_ = 375, *p*_*E*_ = *p*_*I*_ = 0.1 and *g*_*E*_ = *g*_*I*_ = 1.5. Its performance on the standard test signal was 80.5%

## Notes

### Competing Interest Statement

H. Wagner is shareholder in Predimo GmbH.
The other authors have no competing interests that could have influenced the results of this paper.

### Summary of Updates

Revision based on reviewer feedback: (1) minor rewrite to improve clarity of the used terminology, (2) added more literature on muscle synergies and neuromechanical models, (3) added a figure showing neural activity and corresponding principal components, (4) tested the networks on a morphing task, and (5) performed an additional experiment to test the generalizability of the neural network models to other speeds.

